# Classification of Human Chronotype Based on fMRI Network-Based Statistics

**DOI:** 10.1101/2022.08.25.505246

**Authors:** Sophie L. Mason, Leandro Junges, Wessel Woldman, Elise R. Facer-Childs, Brunno M. de Campos, Andrew P. Bagshaw, John R. Terry

## Abstract

Chronotype - the relationship between the internal circadian physiology of an individual and the external 24-hour light-dark cycle - is increasingly implicated in mental health and cognition. Individuals presenting with a late chronotype have an increased likelihood of developing depression, and can display reduced cognitive performance during the societal 9-5 day. However, the interplay between physiological rhythms and the brain networks that underpin cognition and mental health are not well understood. To address this issue, we use resting state fMRI collected from 16 people with an early chronotype and 22 people with a late chronotype to study if differentiable information about chronotype is embedded in functional brain networks. We develop a classifier utilising the Network Based-Statistic (NBS) methodology, using rigorous selection criteria to select t-statistic thresholds within the NBS approach. We find significant differences in functional networks measured in early and late chronotypes and describe conditions under which the classifier achieves 97.3% accuracy. Revealing differences in functional brain networks based on extreme chronotype suggests future avenues of research that may ultimately better characterise the relationship between internal physiology, external perturbations, brain networks and disease.

## 1 Introduction

Almost all bodily functions depend fundamentally on oscillations. A diversity of biological clocks tightly interconnect to control processes over time scales of hours such as sleep stages or body temperature through to days and months, for example, hormone release or the female menstrual cycle (Kondratova and Kondratov 2012).

Processes that oscillate with a period of around a day are called *circadian*. These include neurobehavioral (i.e. attention or mood), hormonal (i.e. melatonin or cortisol secretion) and physiological (i.e. heart rate or body temperature) (A. Wirz-Justice 2007). Circadian rhythms in the context of sleep refer to the naturally occurring oscillatory nature of a human’s high and low sleep propensity (Borbély and Achermann 1999). However, the phase relationship between the internal circadian rhythm and the external clock time can differ markedly between individuals, contributing to a classification of people according to their chronotypes (Blautzik et al. 2013). Chronotype classification will fall across a spectrum, with early circadian phenotypes (ECPs) and late circadian phenotypes (LCPs) sitting at the two extremes. These two extreme phenotypes are often colloquially referred to as ‘Morning Larks’ and ‘Night Owls’. Upon removing external social obligations, the sleep of ECPs and LCPs will show little difference - except for the phase between circadian rhythm and clock time. For example, driven by the endogenous circadian rhythm of the individual, extreme ECPs may wake up at the same time that extreme LCPs are falling to sleep (Roenneberg, Anna Wirz-Justice, and Merrow 2003).

The Munich Chronotype Questionnaire (MCTQ), first introduced in (Roenneberg, Anna Wirz-Justice, and Merrow 2003), is a standard tool for chronotype classification alongside actigraphy data and finding peaks in cortisol and melatonin concentrations from saliva samples (E. R. Facer-Childs 2018). These methods permit classification based upon physiological markers; however, they do not provide an insight into the underlying processes that lead to circadian rhythmicity and how these oscillations ultimately impact upon health and brain function.

Chronotype is a known risk factor for a variety of common health conditions. In particular, late chronotypes are associated with an increased risk of cancer, type 2 diabetes, as well as increased BMI and obesity (Hug et al. 2019). Further, late chronotype is a known risk factor for the development of depression (Levandovski et al. 2011) as well as being predictive of variability in cognitive outcomes across the day (E. R. Facer-Childs 2018). The role of functional brain networks in supporting healthy brain function and the disruptions that lead to impaired performance and disease are increasingly understood in many fields such as depression (Y. Liu et al. 2020) as well as cognition both in terms of cognitive architecture (Petersen and Sporns 2015) as well as cognitive aging (Terry, Anderson, and J. A. Horne 2004, Hausman et al. 2020). Consequently, an increased understanding of the relationship between chronotype and functional brain networks could provide insight into the increased risk of mental health and cognitive outcomes, the reason for the disparity of such impacts along the chronotype spectrum, and therefore areas upon which to focus future research. In addition, a greater understanding of how chronotype impacts the brain’s functional network could provide support for practical changes in society, such as the school day starting later for adolescent children (Adolescent Sleep Working Group And Committee On Adolescence And Council On School Health 2014) or greater flexibility in workplace hours (Vetter et al. 2015).

A common technique to investigate how macroscale neurological processes occur or manifest in the brain is fMRI (functional Magnetic Resonance Imaging). This technique permits regions of interest (ROIs) in the brain to be represented as nodes, with connections (edges) between them determined by statistical interdependencies between their time-series (so-called functional connectivity (FC))(Friston 2011). The FC between a given voxel and all other voxels is called a seed-based approach (Elise R. Facer-Childs, Brunno M. Campos, et al. 2019; Fafrowicz et al. 2019) while representing the brain as a functional network (FN) permits tools and techniques from mathematics, such as graph theory (Farahani, Fafrowicz, Karwowski, Bohaterewicz, et al. 2021; H. Liu et al. 2014) to be utilised. Motivated by the success of seed-based and graph metric pipelines within areas such as time of day effects (Hodkinson et al. 2014), as well as chronic acute (Farahani, Fafrowicz, Karwowski, Douglas, et al. 2019) and prolonged sleep deprivation (H. Liu et al. 2014), some limited studies exploring the role of chronobiology on functional networks have been undertaken. For example, Facer-Childs (Elise R. Facer-Childs, Brunno M. Campos, et al. 2019) seeded in the medial Prefrontal Cortex (mPFC) and Posterior Cingulate Cortex (PCC) within the Default Mode Network (DMN). For both seeds, using the contrast ECP>LCP, the respective clusters were predictive of attentional performance as measured through a psychomotor vigilance test. In addition, resting state FC (rs-FC) recorded from mPFC was predictive of Stroop performance, and PCC was predictive of subjective sleepiness measured using the Karolinska Sleepiness Scale (KSS). Facer-Childs also completed a similar analysis within the Motor Network (MN) system (Elise R. Facer-Childs, Brunno M. de Campos, et al. 2021), where MN rs-FC was shown to contribute to individual variability in motor performance. On the other hand, (Fafrowicz et al. 2019) seeded in thirty-six regions throughout the brain. Correlation of the resulting clusters’ FC to time of day, chronotype, and time of day x chronotype was considered. The effect of chronotype manifested itself differently in these studies: chronotype alongside rs-fMRI was successfully used as a predictor of cognition (Elise R. Facer-Childs, Brunno M. Campos, et al. 2019), whereas (Fafrowicz et al. 2019) failed to find a significant difference between extreme chronotypes with rs-fMRI and instead only found significant time of day effects.

On the other hand, Farahani et al (Farahani, Fafrowicz, Karwowski, Bohaterewicz, et al. 2021) chose a more traditional approach of creating individual FNs before binarising the networks at a range of density-based thresholding values. Graph metrics were then calculated using the binarised networks before being considered for group-level differences between ECPs and LCPs in the morning and evening. Five global measures ranging from network resilience to functional integration and functional segregation as well as combined effects such as small-world index were considered, alongside four local measures. In all cases, graph metrics did not significantly differ based upon chronotype. However, significant time of day effects were once again seen when comparing graph metrics associated with different scans in the morning and evening for the small-world index, and within specific nodes when calculating degree centrality and betweenness centrality.

The limited research and conflicting findings on the role of chronotype on functional networks derived from fMRI data is suggestive of chronotype having relatively subtle effects on functional brain networks, that cannot easily be detected using traditional FC pipelines. An alternative approach is Network Based Statistic (NBS), which is a graph-based method, that aims to find a subnetwork consisting of edges showing high differentiation levels between the functional networks of two groups (typically a control and patient cohort) - called a *dysconnected network*. The term dysconnected network is not to be confused with *disconnected network* a word commonly used within graph theory to refer to a network in which two nodes have no path between them. The original application of NBS explored differences between the functional networks of people with schizophrenia and controls (Zalesky, Fornito, and Bullmore 2010). Since then, NBS has been explored in a range of neurological contexts from the effects of habitual coffee drinking (Magalhães et al. 2021) to considering how age and intelligence (DeSerisy et al. 2021) or symptoms of ADHD (Cocchi et al. 2012) can be correlated with the FC in dysconnected edges found from NBS. A recent extension of NBS (NBS-Predict) has also been published (Serin et al. 2021). It uses a training set to select relevant features from NBS networks for the classification of test subjects. In contrast, our classifier evaluates whether an ECP or LCP label of the test subject leads to greater differentiation between the two groups in a group-level comparison. The use of the test subject therefore differs between the two classification approaches.

The methodology underpinning the NBS approach is described in detail in (Zalesky, Fornito, and Bullmore 2010) and is published alongside a freely available NBS toolbox for MATLAB (MathWorks, USA). Alongside the recent extension NBS-Predict for the classification of test subjects (Serin et al. 2021). For completeness, a brief overview of NBS in the context of chronotype is provided below.

To consider whether NBS may be better suited than seed-based or graph metric pipelines for differentiation between ECPs and LCPs we introduce a classifier to assess the ability of NBS-derived dysconnected networks for classifying individuals as ECP, LCP or unclear. Within NBS, a t-statistic thresholding step is required, therefore a key aim is to create a classifier which includes an objective t-statistic threshold selection process. This avoids multiple comparison issues that may arise from varying the t-statistic threshold over a parameter range. The t-statistic threshold selection criteria is based on the assumption of connectedness for the undirected dysconnected networks. In particular, t-statistic thresholds which create minimum connected components (MCCs) are utilised, therefore ensuring all ROIs are present with an undetermined number of edges. With no prior knowledge of the subjects this accounts for the potential uniqueness in an individual’s subnetwork of importance by assuming all ROIs provide information and must therefore form part of the subnetwork.

Broadly, the classifier uses NBS to create and compare the dysconnected networks resulting from labelling a test subject as first an ECP and then an LCP. We compare the significance of these dysconnected networks at specific t-statistic thresholds, alongside their size (number of edges), to classify the test subject - significance and larger networks are taken as an indication of correct labelling.

Additional analyses were performed examining time of day effects as well as the stability of the classifier on subsets of the dataset created from leaving out one subject at a time. Furthermore, an investigation into the reasons why the removal of certain subjects resulted in large changes in accuracy and possible ways to mitigate this effect were considered.

## 2 Materials and Methods

The chronotype dataset analysed in this study was first presented by (Elise R. Facer-Childs, Brunno M. Campos, et al. 2019), which can be referred to for a detailed description of the data acquisition and preprocessing methodology. Ethical approval was provided by the University of Birmingham Research Ethics Committee, before any data acquisition started. Participants had at least 48 hours before the start of the study to read over information sheets and consent forms, and they were free to end their participation at any time. In addition, the University of Birmingham’s Advisory Group on the Control of Biological Hazards approved the COSHH risk assessments and biological assessment forms that were completed. All the data and samples were given by participants voluntarily and were fully anonymised.

### 2.1 Participants

38 participants were enrolled into the study, with an average age of 22.7 ± 4.2 years (mean ± standard deviation), of whom 24 were female. Exclusion criteria included 1) prior diagnosis of any sleep, neurological or psychiatric disorders 2) use of medication that would knowingly affect sleep 3) an intermediate chronotype classification from the MCTQ. For two weeks prior to the scanning sessions, participants were instructed to follow their preferred sleep pattern, with no restrictions imposed by the study except for two hours before the scanning sessions when alcohol and caffeine consumption, as well as exercise, were prohibited.

16 subjects were classified as ECPs (age 24.7 ± 4.6 years, 9 females) and 22 as LCPs (age 21.3 ± 3.3 years, 15 female). Chronotypes were determined by the outcome of the MCTQ and was further validated by the analysis of Dim Light Melatonin Onset (DLMO) and Cortisol Awakening Response (CAR) times as well as sleep onset and wake up times measured from actigraphy data (Elise R. Facer-Childs, Middleton, et al. 2020). Across all 5 categories, the difference between the groups was significant (Elise R. Facer-Childs, Brunno M. Campos, et al. 2019).

Participants attended three scanning sessions at the local times of 14: 00, 20: 00 and 08: 00 the following day. One ECP (Subject 11) had their 14: 00 rs-fMRI scan excluded due to excessive movement.

### 2.2 Data Acquisition

A Philips Achieva 3 Tesla MRI scanner with a 32-channel head coil was used to collect the imaging data for all participants. This involved a 5-minute T1-weighted scan to create a standard high-resolution anatomical image of the brain (1mm isotropic voxels) before a 15-minute eyes-open resting-state scan was obtained. The scans captured the entire brain using gradient echo echo-planer imaging oriented parallel to the AC-PC line with the following parameters: 450 volumes, TR=2000ms, TE = 35ms, flip angle = 80°, 3 × 3 × 4mm^3^ voxels. Respiratory and cardiac fluctuations were monitored using equipment also provided by Philips.

### 2.3 Data Preprocessing

Pre-processing was completed using UF^2^C (User-Friendly Functional Connectivity) introduced in (Brunno Machado de Campos et al. 2016). This is a toolbox written in MATLAB, which relies upon SPM12 (Penny et al. 2007) and PhysIO (Kasper et al. 2017), which are needed for physiological noise correction. Preprocessing involved standardised steps including re-orientation of scans, motion correction, co-registration, spatially normalising scans to Montreal Neurological Institute (MNI) space, spatially smoothing the data with a 6mm Gaussian kernel, high pass temporal filtering (> 0.01 Hz) and correction for physiological noise, (Elise R. Facer-Childs, Brunno M. Campos, et al. 2019) can be referred to for more additional details.

### 2.4 Functional Network Construction

The brain was parcellated into 70 functional ROIs previously used in (Brunno Machado de Campos et al. 2016) and based upon the 90 functional ROIs used by the Stanford Find Lab presented in Shirer et al. (Shirer et al. 2012). This excluded ROIs in the cerebellum due to incomplete coverage. Information about the ROIs used in the study, including MNI coordinates, is provided in Table 12 in Appendix A. The average timeseries for all voxels within each ROI was then extracted before each timeseries was standardised to mean 0 and standard deviation 1.

**Table 1:**
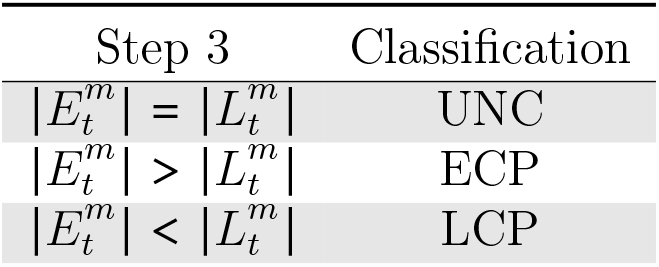
Step Three Streamlined: NBS Classifier. The third step of the NBS classifier, which is needed only if 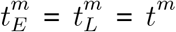 and the two dysconnected networks 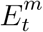 and 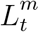 are significant. This step compares the number of edges at threshold *t^m^* when Subject *m* is labelled an ECP and then an LCP.

**Table 2:** Table showing the breakdown of classifications at each step. The number of ECPs and LCPs that are classified at each step of the classification pipeline is presented in the table as well as the accuracy of the classification. The † denotes Subject 36 who was incorrectly labelled at step three of the classifier.

|  | ECP | LCP | Total |
| --- | --- | --- | --- |
| Step one | 10 | 14 | 24 |
| Step two | 1 | 1 | 2 |
| Step three | 5 | 7<sup>†</sup> | 12 |
| Accuracy % | 100 | 95.5 | 97.3 |

**Table 3:** The labels assigned by the classifier for the different scanning sessions and contrasts. Here Subjects 1-16 are ECPs while 17-38 are LCPs, as labelled using non-imaging data.

| Subject | Afternoon<br>ECP<LCP | Afternoon<br>ECP>LCP | Evening<br>ECP<LCP | Evening<br>ECP>LCP | Morning<br>ECP<LCP | Morning<br>ECP>LCP |
| --- | --- | --- | --- | --- | --- | --- |
| 1 | UNC | UNC | UNC | ECP | UNC | UNC |
| 2 | UNC | UNC | UNC | ECP | LCP | UNC |
| 3 | UNC | UNC | UNC | ECP | UNC | UNC |
| 4 | UNC | UNC | UNC | ECP | UNC | UNC |
| 5 | LCP | UNC | UNC | ECP | UNC | UNC |
| 6 | LCP | UNC | UNC | ECP | UNC | UNC |
| 7 | LCP | UNC | UNC | ECP | UNC | UNC |
| 8 | UNC | UNC | UNC | ECP | UNC | UNC |
| 9 | UNC | UNC | UNC | ECP | UNC | UNC |
| 10 | LCP | UNC | UNC | ECP | LCP | UNC |
| 11 | NA | NA | UNC | ECP | UNC | UNC |
| 12 | UNC | UNC | UNC | ECP | UNC | UNC |
| 13 | UNC | UNC | UNC | ECP | LCP | UNC |
| 14 | UNC | UNC | UNC | ECP | UNC | UNC |
| 15 | UNC | UNC | UNC | ECP | UNC | UNC |
| 16 | LCP | UNC | UNC | ECP | UNC | UNC |
| 17 | UNC | UNC | UNC | LCP | UNC | UNC |
| 18 | UNC | UNC | UNC | LCP | UNC | UNC |
| 19 | UNC | UNC | UNC | LCP | ECP | UNC |
| 20 | UNC | UNC | UNC | LCP | ECP | UNC |
| 21 | UNC | UNC | UNC | LCP | UNC | UNC |
| 22 | UNC | ECP | UNC | LCP | UNC | UNC |
| 23 | ECP | UNC | UNC | LCP | UNC | UNC |
| 24 | UNC | UNC | UNC | LCP | UNC | UNC |
| 25 | UNC | UNC | UNC | LCP | UNC | UNC |
| 26 | ECP | UNC | UNC | LCP | ECP | UNC |
| 27 | UNC | UNC | UNC | LCP | ECP | UNC |
| 28 | UNC | UNC | UNC | LCP | UNC | UNC |
| 29 | UNC | UNC | UNC | LCP | UNC | UNC |
| 30 | UNC | UNC | UNC | LCP | UNC | UNC |
| 31 | ECP | UNC | UNC | LCP | UNC | UNC |
| 32 | UNC | UNC | UNC | LCP | UNC | UNC |
| 33 | UNC | UNC | UNC | LCP | UNC | UNC |
| 34 | UNC | UNC | UNC | LCP | UNC | UNC |
| 35 | UNC | UNC | UNC | LCP | UNC | UNC |
| 36 | ECP | UNC | UNC | ECP | ECP | UNC |
| 37 | ECP | UNC | UNC | LCP | ECP | UNC |
| 38 | ECP | UNC | UNC | LCP | UNC | UNC |
| Accuracy(%) | 0 | 0 | 0 | 97.3 | 0 | 0 |
| Percentage Classified<br>ECP or LCP (%) | 29.7 | 2.7 | 0 | 100 | 23.7 | 0 |

**Table 4:**
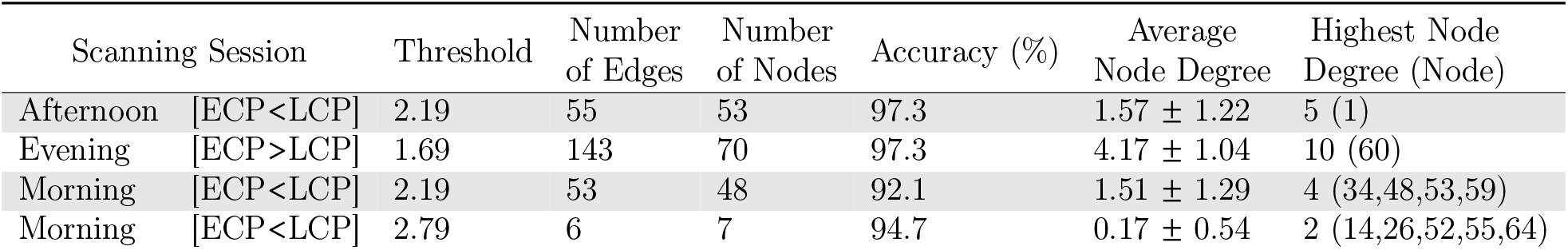
Table Showing Key Topological Features of Four dysconnected Networks.

| Scanning Session |  | Threshold | Number of Edges | Number of Nodes | Accuracy (%) | Average Node Degree | Highest Node Degree (Node) |
| --- | --- | --- | --- | --- | --- | --- | --- |
| Afternoon | [ECP<LCP] | 2.19 | 55 | 53 | 97.3 | $1.57 \pm 1.22$ | 5 (1) |
| Evening | [ECP>LCP] | 1.69 | 143 | 70 | 97.3 | $4.17 \pm 1.04$ | 10 (60) |
| Morning | [ECP<LCP] | 2.19 | 53 | 48 | 92.1 | $1.51 \pm 1.29$ | 4 (34,48,53,59) |
| Morning | [ECP<LCP] | 2.79 | 6 | 7 | 94.7 | $0.17 \pm 0.54$ | 2 (14,26,52,55,64) |

**Table 5:**
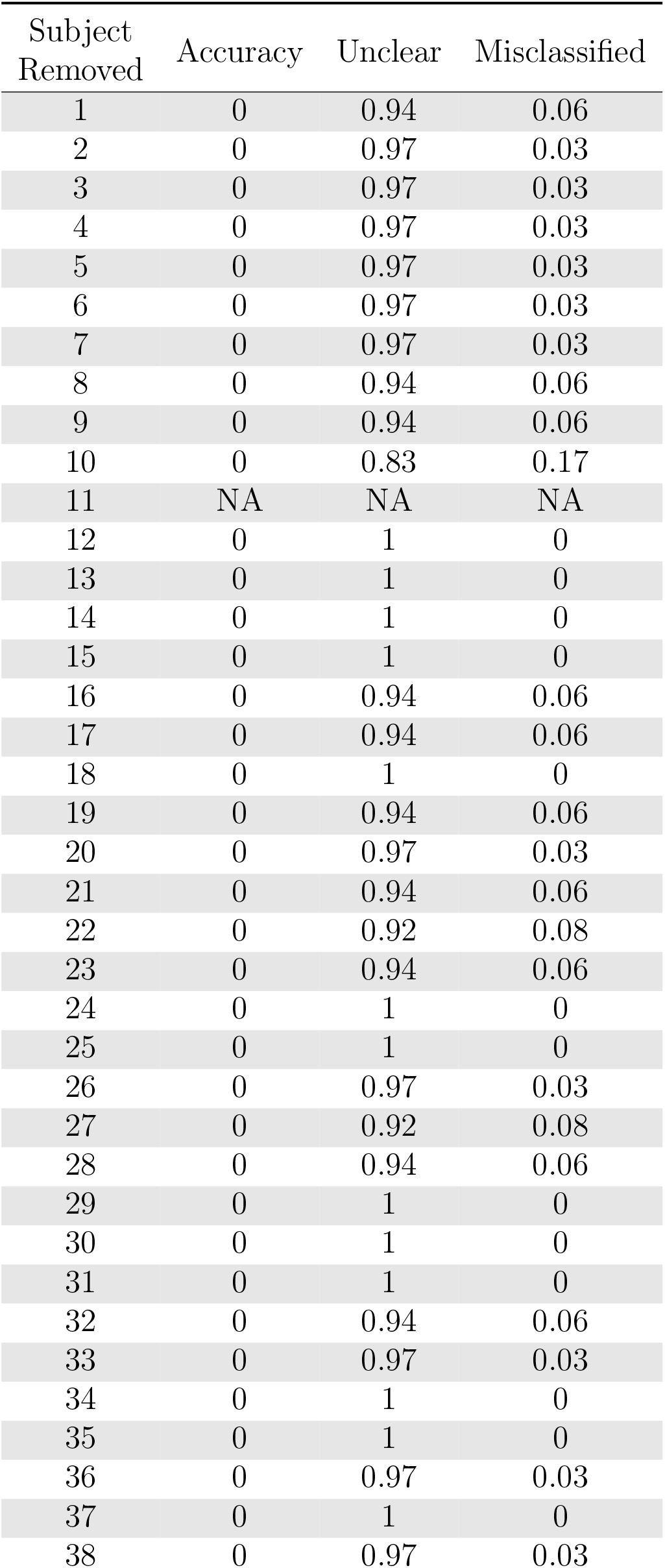
Afternoon [ECP>LCP] Results: Partial Datasets. The percentage of correct, unclear and incorrect classifications when one subject was removed.

| Subject<br>Removed | Accuracy | Unclear | Misclassified |
| --- | --- | --- | --- |
| 1 | 0 | 0.94 | 0.06 |
| 2 | 0 | 0.97 | 0.03 |
| 3 | 0 | 0.97 | 0.03 |
| 4 | 0 | 0.97 | 0.03 |
| 5 | 0 | 0.97 | 0.03 |
| 6 | 0 | 0.97 | 0.03 |
| 7 | 0 | 0.97 | 0.03 |
| 8 | 0 | 0.94 | 0.06 |
| 9 | 0 | 0.94 | 0.06 |
| 10 | 0 | 0.83 | 0.17 |
| 11 | NA | NA | NA |
| 12 | 0 | 1 | 0 |
| 13 | 0 | 1 | 0 |
| 14 | 0 | 1 | 0 |
| 15 | 0 | 1 | 0 |
| 16 | 0 | 0.94 | 0.06 |
| 17 | 0 | 0.94 | 0.06 |
| 18 | 0 | 1 | 0 |
| 19 | 0 | 0.94 | 0.06 |
| 20 | 0 | 0.97 | 0.03 |
| 21 | 0 | 0.94 | 0.06 |
| 22 | 0 | 0.92 | 0.08 |
| 23 | 0 | 0.94 | 0.06 |
| 24 | 0 | 1 | 0 |
| 25 | 0 | 1 | 0 |
| 26 | 0 | 0.97 | 0.03 |
| 27 | 0 | 0.92 | 0.08 |
| 28 | 0 | 0.94 | 0.06 |
| 29 | 0 | 1 | 0 |
| 30 | 0 | 1 | 0 |
| 31 | 0 | 1 | 0 |
| 32 | 0 | 0.94 | 0.06 |
| 33 | 0 | 0.97 | 0.03 |
| 34 | 0 | 1 | 0 |
| 35 | 0 | 1 | 0 |
| 36 | 0 | 0.97 | 0.03 |
| 37 | 0 | 1 | 0 |
| 38 | 0 | 0.97 | 0.03 |

**Table 6:** Afternoon [ECP<LCP] Results: Partial Datasets. The percentage of correct, unclear and incorrect classifications when one subject was removed

| Subject<br>Removed | Accuracy | Unclear | Misclassified |
| --- | --- | --- | --- |
| 1 | 0 | 0.75 | 0.25 |
| 2 | 0 | 0.83 | 0.17 |
| 3 | 0 | 0.81 | 0.19 |
| 4 | 0 | 0.89 | 0.11 |
| 5 | 0.64 | 0.03 | 0.33 |
| 6 | 0.72 | 0.11 | 0.17 |
| 7 | 0.69 | 0.03 | 0.28 |
| 8 | 0 | 0.78 | 0.22 |
| 9 | 0 | 0.78 | 0.22 |
| 10 | 0.78 | 0.003 | 0.19 |
| 11 | NA | NA | NA |
| 12 | 0 | 0.86 | 0.14 |
| 13 | 0 | 0.81 | 0.19 |
| 14 | 0 | 0.97 | 0.03 |
| 15 | 0 | 0.61 | 0.39 |
| 16 | 0.81 | 0.11 | 0.083 |
| 17 | 0 | 0.92 | 0.08 |
| 18 | 0 | 0.83 | 0.17 |
| 19 | 0 | 0.78 | 0.22 |
| 20 | 0 | 0.75 | 0.25 |
| 21 | 0 | 0.75 | 0.25 |
| 22 | 0 | 0.78 | 0.22 |
| 23 | 0.61 | 0.06 | 0.33 |
| 24 | 0 | 0.72 | 0.28 |
| 25 | 0 | 0.86 | 0.14 |
| 26 | 0.64 | 0 | 0.36 |
| 27 | 0 | 0.89 | 0.11 |
| 28 | 0 | 0.75 | 0.25 |
| 29 | 0 | 0.78 | 0.22 |
| 30 | 0 | 0.86 | 0.14 |
| 31 | 0 | 0.69 | 0.31 |
| 32 | 0 | 0.94 | 0.06 |
| 33 | 0 | 0.81 | 0.19 |
| 34 | 0 | 0.94 | 0.06 |
| 35 | 0 | 0.72 | 0.28 |
| 36 | 0 | 0.67 | 0.33 |
| 37 | 0 | 0.67 | 0.33 |
| 38 | 0.39 | 0.28 | 0.33 |

**Table 7:** Evening [ECP>LCP] Results: Partial Datasets. The percentage of correct, unclear and incorrect classifications when one subject was removed

| Subject<br>Removed | Accuracy | Unclear | Misclassified |
| --- | --- | --- | --- |
| 1 | 0.65 | 0.14 | 0.22 |
| 2 | 0.97 | 0.03 | 0 |
| 3 | 0.78 | 0.08 | 0.14 |
| 4 | 0.78 | 0.11 | 0.11 |
| 5 | 0.95 | 0.03 | 0.03 |
| 6 | 0.84 | 0.03 | 0.14 |
| 7 | 0 | 0.76 | 0.24 |
| 8 | 0 | 0.81 | 0.19 |
| 9 | 0 | 0.73 | 0.27 |
| 10 | 0.73 | 0.08 | 0.19 |
| 11 | 0.65 | 0.16 | 0.19 |
| 12 | 0.70 | 0.14 | 0.16 |
| 13 | 0.81 | 0.19 | 0 |
| 14 | 0.81 | 0.08 | 0.11 |
| 15 | 0.95 | 0.05 | 0 |
| 16 | 0 | 0.78 | 0.22 |
| 17 | 0.70 | 0.08 | 0.22 |
| 18 | 0.41 | 0.32 | 0.27 |
| 19 | 0 | 0.78 | 0.22 |
| 20 | 0.86 | 0.05 | 0.08 |
| 21 | 0.95 | 0 | 0.05 |
| 22 | 0.43 | 0.41 | 0.16 |
| 23 | 0.86 | 0.05 | 0.08 |
| 24 | 0.81 | 0.03 | 0.16 |
| 25 | 0 | 0.81 | 0.19 |
| 26 | 0.84 | 0.08 | 0.08 |
| 27 | 0.68 | 0.05 | 0.27 |
| 28 | 0.59 | 0.16 | 0.24 |
| 29 | 0.92 | 0.03 | 0.05 |
| 30 | 0.43 | 0.38 | 0.19 |
| 31 | 0.86 | 0.03 | 0.11 |
| 32 | 0 | 0.78 | 0.22 |
| 33 | 0.76 | 0.11 | 0.14 |
| 34 | 0 | 0.81 | 0.19 |
| 35 | 0.86 | 0.05 | 0.08 |
| 36 | 0.76 | 0.19 | 0.05 |
| 37 | 0.62 | 0.19 | 0.19 |
| 38 | 0.81 | 0.05 | 0.14 |

**Table 8:** Evening [ECP<LCP] Results: Partial Datasets. The percentage of correct, unclear and incorrect classifications when one subject was removed

| Subject<br>Removed | Accuracy | Unclear | Misclassified |
| --- | --- | --- | --- |
| 1 | 0 | 1 | 0 |
| 2 | 0 | 1 | 0 |
| 3 | 0 | 1 | 0 |
| 4 | 0 | 1 | 0 |
| 5 | 0 | 1 | 0 |
| 6 | 0 | 1 | 0 |
| 7 | 0 | 1 | 0 |
| 8 | 0 | 1 | 0 |
| 9 | 0 | 1 | 0 |
| 10 | 0 | 1 | 0 |
| 11 | 0 | 1 | 0 |
| 12 | 0 | 1 | 0 |
| 13 | 0 | 1 | 0 |
| 14 | 0 | 1 | 0 |
| 15 | 0 | 1 | 0 |
| 16 | 0 | 1 | 0 |
| 17 | 0 | 1 | 0 |
| 18 | 0 | 1 | 0 |
| 19 | 0 | 1 | 0 |
| 20 | 0 | 1 | 0 |
| 21 | 0 | 1 | 0 |
| 22 | 0 | 1 | 0 |
| 23 | 0 | 1 | 0 |
| 24 | 0 | 1 | 0 |
| 25 | 0 | 1 | 0 |
| 26 | 0 | 0.97 | 0.03 |
| 27 | 0 | 1 | 0 |
| 28 | 0 | 1 | 0 |
| 29 | 0 | 1 | 0 |
| 30 | 0 | 1 | 0 |
| 31 | 0 | 1 | 0 |
| 32 | 0 | 1 | 0 |
| 33 | 0 | 1 | 0 |
| 34 | 0 | 1 | 0 |
| 35 | 0 | 1 | 0 |
| 36 | 0 | 1 | 0 |
| 37 | 0 | 1 | 0 |
| 38 | 0 | 1 | 0 |

**Table 9:**
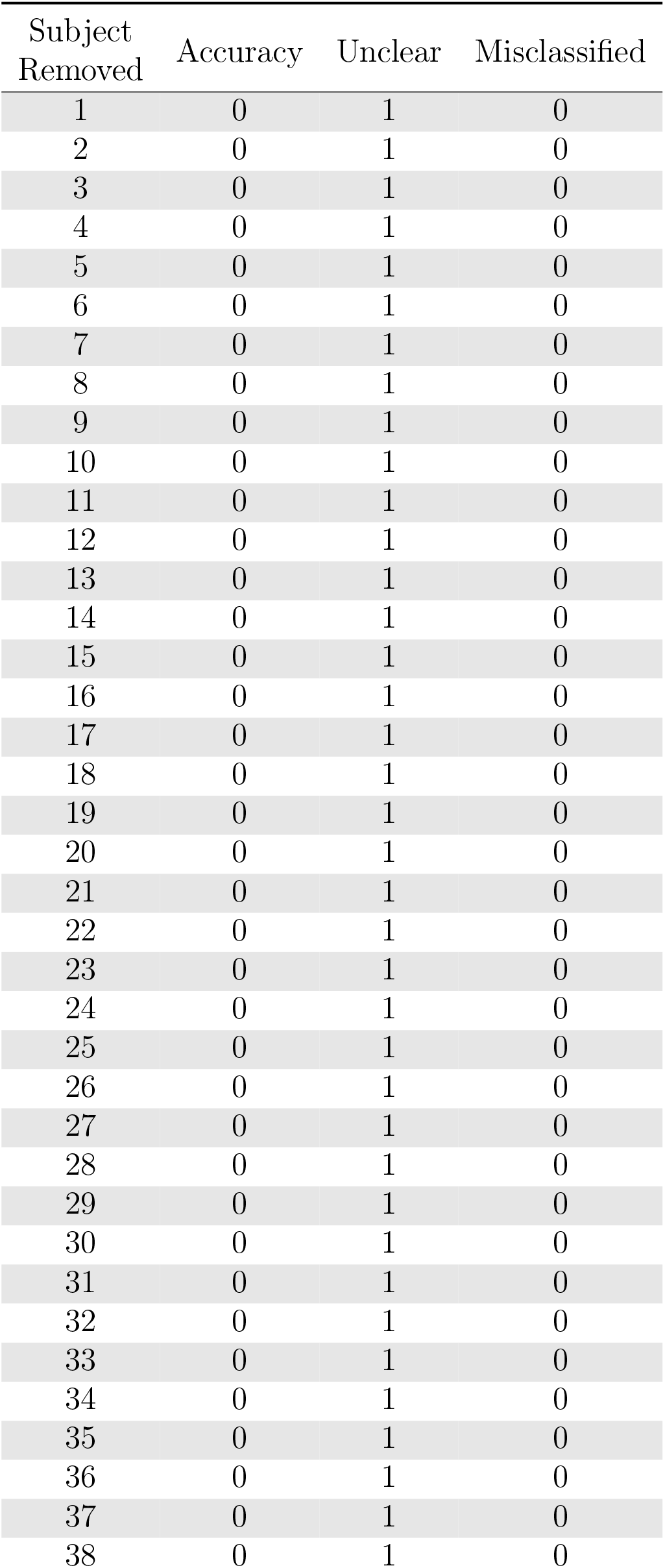
Morning [ECP>LCP] Results: Partial Datasets. The percentage of correct, unclear and incorrect classifications when one subject was removed

| Subject<br>Removed | Accuracy | Unclear | Misclassified |
| --- | --- | --- | --- |
| 1 | 0 | 1 | 0 |
| 2 | 0 | 1 | 0 |
| 3 | 0 | 1 | 0 |
| 4 | 0 | 1 | 0 |
| 5 | 0 | 1 | 0 |
| 6 | 0 | 1 | 0 |
| 7 | 0 | 1 | 0 |
| 8 | 0 | 1 | 0 |
| 9 | 0 | 1 | 0 |
| 10 | 0 | 1 | 0 |
| 11 | 0 | 1 | 0 |
| 12 | 0 | 1 | 0 |
| 13 | 0 | 1 | 0 |
| 14 | 0 | 1 | 0 |
| 15 | 0 | 1 | 0 |
| 16 | 0 | 1 | 0 |
| 17 | 0 | 1 | 0 |
| 18 | 0 | 1 | 0 |
| 19 | 0 | 1 | 0 |
| 20 | 0 | 1 | 0 |
| 21 | 0 | 1 | 0 |
| 22 | 0 | 1 | 0 |
| 23 | 0 | 1 | 0 |
| 24 | 0 | 1 | 0 |
| 25 | 0 | 1 | 0 |
| 26 | 0 | 1 | 0 |
| 27 | 0 | 1 | 0 |
| 28 | 0 | 1 | 0 |
| 29 | 0 | 1 | 0 |
| 30 | 0 | 1 | 0 |
| 31 | 0 | 1 | 0 |
| 32 | 0 | 1 | 0 |
| 33 | 0 | 1 | 0 |
| 34 | 0 | 1 | 0 |
| 35 | 0 | 1 | 0 |
| 36 | 0 | 1 | 0 |
| 37 | 0 | 1 | 0 |
| 38 | 0 | 1 | 0 |

**Table 10:**
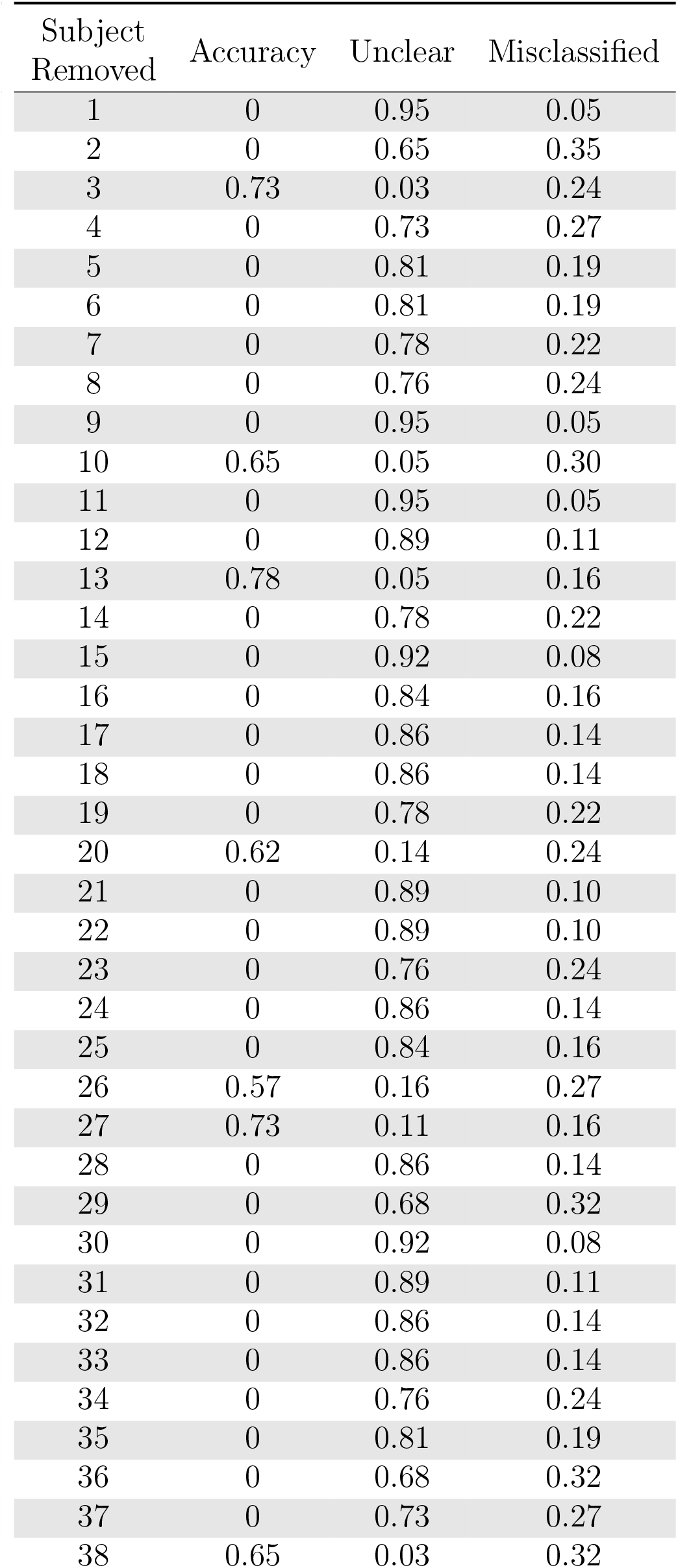
Morning [ECP<LCP] Results: Partial Datasets. The percentage of correct, unclear and incorrect classifications when one subject was removed

| Subject<br>Removed | Accuracy | Unclear | Misclassified |
| --- | --- | --- | --- |
| 1 | 0 | 0.95 | 0.05 |
| 2 | 0 | 0.65 | 0.35 |
| 3 | 0.73 | 0.03 | 0.24 |
| 4 | 0 | 0.73 | 0.27 |
| 5 | 0 | 0.81 | 0.19 |
| 6 | 0 | 0.81 | 0.19 |
| 7 | 0 | 0.78 | 0.22 |
| 8 | 0 | 0.76 | 0.24 |
| 9 | 0 | 0.95 | 0.05 |
| 10 | 0.65 | 0.05 | 0.30 |
| 11 | 0 | 0.95 | 0.05 |
| 12 | 0 | 0.89 | 0.11 |
| 13 | 0.78 | 0.05 | 0.16 |
| 14 | 0 | 0.78 | 0.22 |
| 15 | 0 | 0.92 | 0.08 |
| 16 | 0 | 0.84 | 0.16 |
| 17 | 0 | 0.86 | 0.14 |
| 18 | 0 | 0.86 | 0.14 |
| 19 | 0 | 0.78 | 0.22 |
| 20 | 0.62 | 0.14 | 0.24 |
| 21 | 0 | 0.89 | 0.10 |
| 22 | 0 | 0.89 | 0.10 |
| 23 | 0 | 0.76 | 0.24 |
| 24 | 0 | 0.86 | 0.14 |
| 25 | 0 | 0.84 | 0.16 |
| 26 | 0.57 | 0.16 | 0.27 |
| 27 | 0.73 | 0.11 | 0.16 |
| 28 | 0 | 0.86 | 0.14 |
| 29 | 0 | 0.68 | 0.32 |
| 30 | 0 | 0.92 | 0.08 |
| 31 | 0 | 0.89 | 0.11 |
| 32 | 0 | 0.86 | 0.14 |
| 33 | 0 | 0.86 | 0.14 |
| 34 | 0 | 0.76 | 0.24 |
| 35 | 0 | 0.81 | 0.19 |
| 36 | 0 | 0.68 | 0.32 |
| 37 | 0 | 0.73 | 0.27 |
| 38 | 0.65 | 0.03 | 0.32 |

**Table 11:**
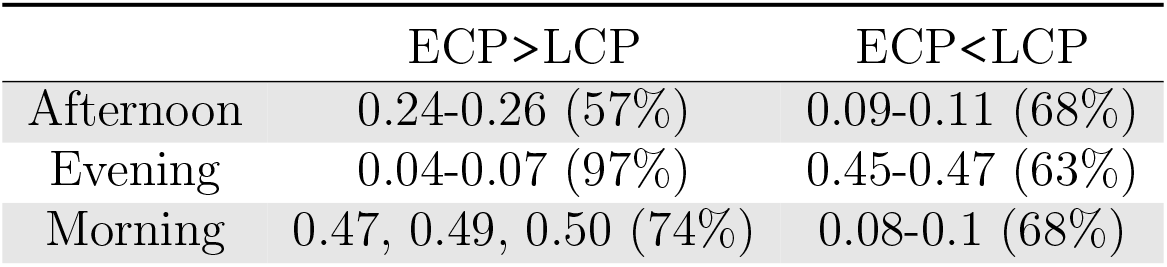
Table showing the values of *α* which give the highest accuracy with the accuracy values given in brackets.

**Table 12:**
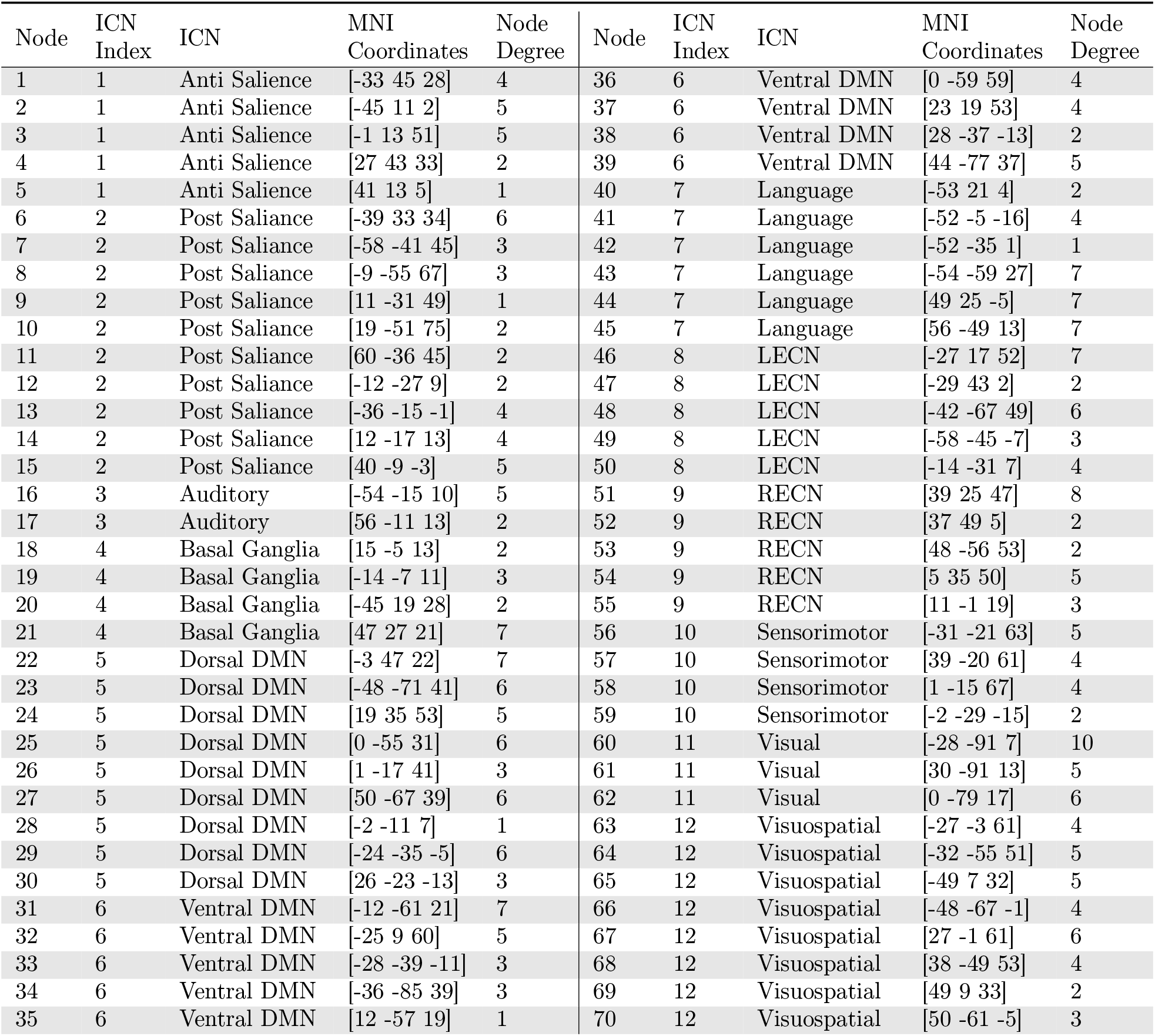
Table giving the nodes, ICNs and MNI Coordinates used for 70 functional connectivity ROIS. In addition, the node degree resulting from the largest connected component for correct labelling for each ROI using the Evening scanning session [ECP>LCP]

| Node | ICN Index | ICN | MNI Coordinates | Node Degree | Node | ICN Index | ICN | MNI Coordinates | Node Degree |
| --- | --- | --- | --- | --- | --- | --- | --- | --- | --- |
| 1 | 1 | Anti Salience | [-33 45 28] | 4 | 36 | 6 | Ventral DMN | [0 -59 59] | 4 |
| 2 | 1 | Anti Salience | [-45 11 2] | 5 | 37 | 6 | Ventral DMN | [23 19 53] | 4 |
| 3 | 1 | Anti Salience | [-1 13 51] | 5 | 38 | 6 | Ventral DMN | [28 -37 -13] | 2 |
| 4 | 1 | Anti Salience | [27 43 33] | 2 | 39 | 6 | Ventral DMN | [44 -77 37] | 5 |
| 5 | 1 | Anti Salience | [41 13 5] | 1 | 40 | 7 | Language | [-53 21 4] | 2 |
| 6 | 2 | Post Salience | [-39 33 34] | 6 | 41 | 7 | Language | [-52 -5 -16] | 4 |
| 7 | 2 | Post Salience | [-58 -41 45] | 3 | 42 | 7 | Language | [-52 -35 1] | 1 |
| 8 | 2 | Post Salience | [-9 -55 67] | 3 | 43 | 7 | Language | [-54 -59 27] | 7 |
| 9 | 2 | Post Salience | [11 -31 49] | 1 | 44 | 7 | Language | [49 25 -5] | 7 |
| 10 | 2 | Post Salience | [19 -51 75] | 2 | 45 | 7 | Language | [56 -49 13] | 7 |
| 11 | 2 | Post Salience | [60 -36 45] | 2 | 46 | 8 | LECN | [-27 17 52] | 7 |
| 12 | 2 | Post Salience | [-12 -27 9] | 2 | 47 | 8 | LECN | [-29 43 2] | 2 |
| 13 | 2 | Post Salience | [-36 -15 -1] | 4 | 48 | 8 | LECN | [-42 -67 49] | 6 |
| 14 | 2 | Post Salience | [12 -17 13] | 4 | 49 | 8 | LECN | [-58 -45 -7] | 3 |
| 15 | 2 | Post Salience | [40 -9 -3] | 5 | 50 | 8 | LECN | [-14 -31 7] | 4 |
| 16 | 3 | Auditory | [-54 -15 10] | 5 | 51 | 9 | RECN | [39 25 47] | 8 |
| 17 | 3 | Auditory | [56 -11 13] | 2 | 52 | 9 | RECN | [37 49 5] | 2 |
| 18 | 4 | Basal Ganglia | [15 -5 13] | 2 | 53 | 9 | RECN | [48 -56 53] | 2 |
| 19 | 4 | Basal Ganglia | [-14 -7 11] | 3 | 54 | 9 | RECN | [5 35 50] | 5 |
| 20 | 4 | Basal Ganglia | [-45 19 28] | 2 | 55 | 9 | RECN | [11 -1 19] | 3 |
| 21 | 4 | Basal Ganglia | [47 27 21] | 7 | 56 | 10 | Sensorimotor | [-31 -21 63] | 5 |
| 22 | 5 | Dorsal DMN | [-3 47 22] | 7 | 57 | 10 | Sensorimotor | [39 -20 61] | 4 |
| 23 | 5 | Dorsal DMN | [-48 -71 41] | 6 | 58 | 10 | Sensorimotor | [1 -15 67] | 4 |
| 24 | 5 | Dorsal DMN | [19 35 53] | 5 | 59 | 10 | Sensorimotor | [-2 -29 -15] | 2 |
| 25 | 5 | Dorsal DMN | [0 -55 31] | 6 | 60 | 11 | Visual | [-28 -91 7] | 10 |
| 26 | 5 | Dorsal DMN | [1 -17 41] | 3 | 61 | 11 | Visual | [30 -91 13] | 5 |
| 27 | 5 | Dorsal DMN | [50 -67 39] | 6 | 62 | 11 | Visual | [0 -79 17] | 6 |
| 28 | 5 | Dorsal DMN | [-2 -11 7] | 1 | 63 | 12 | Visuospatial | [-27 -3 61] | 4 |
| 29 | 5 | Dorsal DMN | [-24 -35 -5] | 6 | 64 | 12 | Visuospatial | [-32 -55 51] | 5 |
| 30 | 5 | Dorsal DMN | [26 -23 -13] | 3 | 65 | 12 | Visuospatial | [-49 7 32] | 5 |
| 31 | 6 | Ventral DMN | [-12 -61 21] | 7 | 66 | 12 | Visuospatial | [-48 -67 -1] | 4 |
| 32 | 6 | Ventral DMN | [-25 9 60] | 5 | 67 | 12 | Visuospatial | [27 -1 61] | 6 |
| 33 | 6 | Ventral DMN | [-28 -39 -11] | 3 | 68 | 12 | Visuospatial | [38 -49 53] | 4 |
| 34 | 6 | Ventral DMN | [-36 -85 39] | 3 | 69 | 12 | Visuospatial | [49 9 33] | 2 |
| 35 | 6 | Ventral DMN | [12 -57 19] | 1 | 70 | 12 | Visuospatial | [50 -61 -5] | 3 |

FNs were then calculated for each subject using Tikhonov partial correlation. A symmetric weighted adjacency matrix with a size 70 × 70 was created for each scan where edge weights are in the range [−1, 1]. Note that all values in the leading diagonal are set to NaN as self-links are not interpretable under partial correlation. Partial correlation was chosen to reduce the contribution from indirect connections (Marrelec, Krainik, et al. 2006; Marrelec, Kim, et al. 2009) as well as its superior performance at replicating ground truth networks considered in simulated data studies (Smith et al. 2011; H. E. Wang et al. 2014; Y. Wang et al. 2016). Further, (Pervaiz et al. 2020) did an extensive review of partial correlation as well as various regularisation techniques, and concluded Tikhonov partial correlation as a recommended method. The regularisation technique requires adding a scaled identity matrix to the covariance matrix of each subject before partial correlation is calculated using the inverse of the (regularised) covariance matrix, known as the (Tikhonov) precision matrix. The scalar λ, known as the regularisation parameter, was optimised by minimising the difference between the average precision matrix across all participants and the Tikhonov precision matrix of each individual participant. This optimisation was completed by summing over all participants the element-wise difference between the average and individual precision matrix before squaring each element. Since the resulting matrix is symmetric, the square root of the sum of the upper triangular elements was calculated. Through considering a range of values of the parameter λ, the choice which minimises this square root sum is considered optimal. For this study, using data from all three scanning sessions λ = 0.0259 was found to be optimal. The regularisation λ = 0.0259 was used for the creation of all Tikhonov partial correlation matrices throughout the paper. Further details on the use of Tikhonov partial correlation, as well as the calculation of the regularisation parameter are presented in Appendix B.

### 2.5 Network-Based Statistics

After calculating FNs for all subjects, we used NBS to determine dysconnected networks. The group level difference in edge weights for every edge in the Tikhonov partial correlation functional networks across subjects was determined using a t-test under a specific contrast - either ECP>LCP or ECP<LCP. In this case each element represents the difference in the mean edge weights between the two groups resulting in a symmetric matrix whose elements are the output of a two-sample one-sided t-test. The t-statistic matrix was then thresholded with the largest connected component of suprathreshold edges, called a dysconnected network, selected as the subnetwork of interest. Here the dysconnected network is a subnetwork of edges that show the highest difference in FC between the two phenotypes. The significance of a dysconnected network was determined using non-parametric permutation testing to create a familywise error (FWE) corrected p-value. This test compares the intensity (the total weight of the edges) of the connected component to a null distribution of connected component intensities created by randomising the group to which each participant is assigned. The p-value was then calculated as the percentage of random permutations whose intensity is larger than that of the measured dysconnected network.

### 2.6 NBS Threshold Selection

Thresholding is a key step within the traditional NBS methodology. Different t-statistic threshold choices can have a critical impact on network properties such as the number of nodes and edges, and therefore impact the properties of the dysconnected network. When seeking to label each test subject as an ECP and subsequently an LCP, a t-statistic threshold is required for each labelling. The choice of the t-statistic threshold in the original NBS literature is somewhat arbitrary, with no rigorous method suggested. Here we suggest a process for determining these t-statistic thresholds, which is as follows:

For each individual test subject *m* ∈ {1, 2,…, 38}, label the test subject an ECP while letting the training set retain their chronotype label (determined by MCTQ, saliva samples and actigraphy data). Using the NBS pipeline calculate the 70 × 70 t-statistic matrix across all subjects using the contrast ECP>LCP or ECP<LCP. Each element represents the difference in the mean edge weights between the two groups of extreme chronotypes when the test subject has been artificially placed in the ECP group. Then set 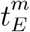 as the highest t-statistic threshold such that the network of suprathreshold edges is connected. This value 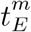 is known as the percolation threshold. Similarly, repeat the steps above with the test subject m labelled an LCP to find the percolation threshold for the LCP labelling, setting 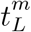 to this value.

The importance of setting t-statistic thresholds 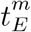 and 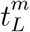 as percolation thresholds is directly linked to the creation of minimum connected components (MCCs), which were first presented and fully explained in (Vijayalakshmi et al. 2015). Effectively, the MCC is a pragmatic balance between the sparsity and density of a network. Interestingly, (Vijayalakshmi et al. 2015) found the MCC to be sensitive to subtle changes in FNs resulting from changes to cognitive load in EEG recordings, which were difficult to detect using other methods.

### 2.7 The Classifier

To determine whether an individual is an ECP or an LCP, we developed a classifier that considers whether the individual fits best within a family of known ECPs or LCPs. The main steps are summarised in Figure 1.

**Figure 1:**
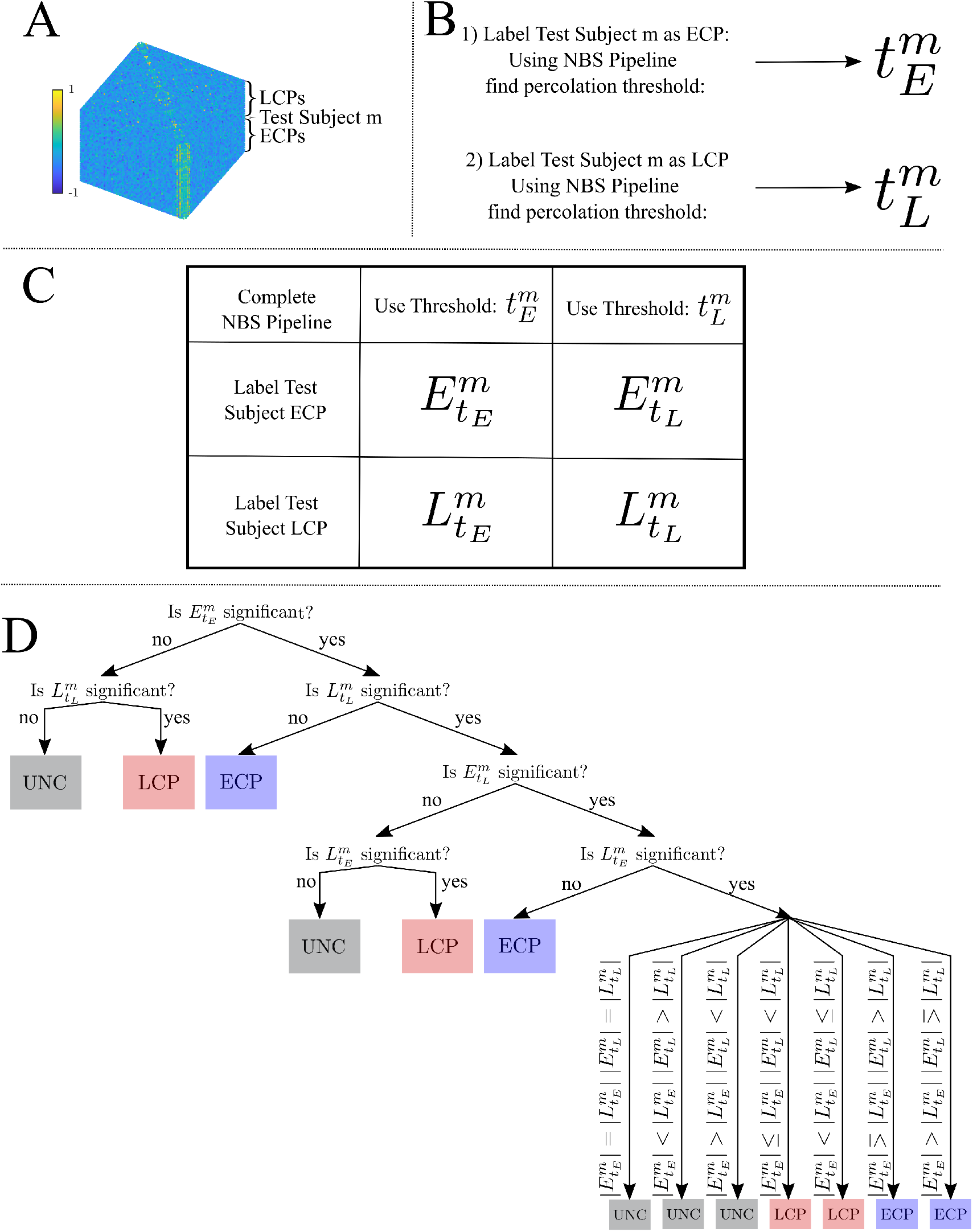
Summary Pipeline for the Three Step Classifier. **(A)** All subjects Tikhonov Partial Correlation matrices (70 × 70 × number of subjects) where test Subject *m* has a label unknown to the classifier. **(B)** Using the NBS pipeline to find the threshold of least redundancy when the test subject has been labelled an ECP and then an LCP. **C)** The four dysconnected networks which are created from the two thresholds and two labellings. **(D)** The three steps of the classifier to classify Subject *m* as an ECP, an LCP or unclear.

Let 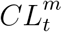 denote the dysconnected network created by assigning subject *m* the chronotype label (*CL*) at t-statistic threshold *t*, calculated using the NBS toolbox (Zalesky, Fornito, and Bullmore 2010). For the two possible chronotype labellings of the test subject and the two t-statistic thresholding values 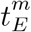 and 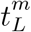 four subnetworks are created: 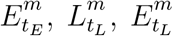 and 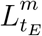. Each subnetwork will have its own FWE-controlled p-value where *N* = 5000 random permutations of class labels were used to create a null distribution. Here *p* < 0.05 is considered to be significant, and intensity was used when calculating the *p*-value. Intensity was used because the NBS reference manual (Zalesky, Fornito, Cocchi, et al. 2012) suggests calculating the FWE *p*-value using intensity rather than the number of edges is beneficial for detecting subtle (distributed but sparse) effects throughout the network, rather than focal effects within a specific component of the network. From here, the classifier was constructed as follows:

Step one is to consider the significance of 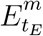 and 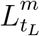. If only one of these is significant, the classification of subject *m* is the significant chronotype label *CL*. If neither are significant the classification of subject *m* is unclear. If both are deemed to have a significant size compared to a null distribution then further steps are needed.

Step two is to consider the significance of the two remaining percolation thresholds for each *CL*: 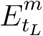 and 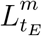. In this case, if only one of these is significant, the classification of subject *m* is the significant chronotype label *CL*. If neither are significant then the classification of subject m is unclear. If both are deemed to have a significant intensity compared to a null distribution, one further step is required.

Step three is to consider the number of edges |*CL_t_*| of the dysconnected networks in t-statistic threshold pairs. For the four significant subnetworks 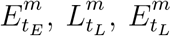 and 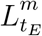, the chronotype label with the higher number of edges indicates the label that the classifier will assign to subject *m* - all comparisons are shown in Figure 1. This is based upon the assumption that for any edge, correctly labelling a subject will result in greater separation between the two groups edge weights, which should be reflected in higher t-statistic values. Therefore, at a specific t-statistic threshold, more edges should survive the thresholding step when correctly labelled in contrast to being incorrectly labelled.

Note that for the accuracy levels presented a classification of unclear is treated as separate to correctly classified and misclassified. Therefore, an unclear classification will not increase the accuracy level.

### 2.8 Varying the t-statistic Threshold

A method for selecting the threshold values at which to threshold the t-statistic matrix has already been outlined in Section 2.6. However, the threshold selection process was reliant on the assumption that all ROIs provide differential information. To understand the impact of this assumption we also decided to range the t-statistic thresholding parameter from 0 to 4.5 in increments of 0.01. Therefore, only one t-statistic threshold is selected for both chronotype labellings.

As the t-statistic threshold range extended beyond the percolation threshold some dysconnected networks did not include all ROIs. Therefore, for certain t-statistic thresholds multiple distinct dysconnected networks existed. In this case, we selected the dysconnected network with the smallest *p*-value. When multiple dysconnected networks had the same *p*-value then the dysconnected network with the highest number of edges was selected. If multiple dysconnected networks had the same number of edges then for the purpose of classification they were indistinguishable to the classifier and one was selected arbitrarily.

After selecting the dysconnected networks it can be noted that when 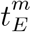 and 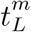 are equal (i.e. ∃ *t^m^* ∈ **R**^+^ s.t. 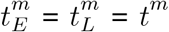) the classifier can be applied as described in Section 2.7, however to reduce redundancy step one can be removed and step three can be streamlined as shown in Table 1. For instance, the case (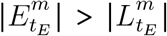 and 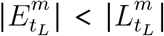) is no longer plausible since 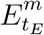 and 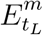 as well as 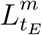 and 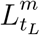 are identical.

Dysconnected networks for select thresholds were then considered and displayed visually using BrainNet Viewer (Xia, J. Wang, and He 2013, http://www.nitrc.org/projects/bnv).

### 2.9 Investigating the Stability of the Classifier

When creating a classifier on small datasets, such as the 38 participants available in this study, a sensible step is to validate the classifier on a similar but independent dataset. This allows you to assess whether the classifier is overfitted to the original data and hence its applicability to different datasets. However, with no access to an alternative existing dataset, surrogate datasets were created using the original data. These partial datasets were created by removing one subject in turn from each scanning session; therefore, creating 113 new datasets, across the three scanning sessions (*n* = 37 Afternoon, *n* = 38 Evening and *n* = 38 Morning).

For each of these new datasets the methodology as outlined above was completed. With accuracy now given as the percentage of correctly labelled subjects calculated from a leave-one-out cross validation (LOOCV) analysis on N-1 subjects. Changing the number of subjects will effect the MCC created using NBS both in terms of the size as well as the specific edges included. This will in turn affect the classification of networks in step one and two through the significance of MCCs changing when a subject is removed, as well as the different number of edges affecting step three of the classifier.

Given the overlap between the partial and full datasets we expect the accuracy levels to be consistent. Indeed, any discrepancies in the accuracy when a single subject is removed could indicate potential problems with the classifier or that some characteristics of the participant’s data are inherently different and influential.

### 2.10 Varying the Threshold for Significance

The dysconnected networks that are created using NBS are considered significant if their *p*-value is less than the significance threshold, *α* = 0.05, such that *p* < 0.05 indicates significance. The threshold of 0.05 is somewhat arbitrary and selected due to the consistent use of this threshold throughout literature. However, the classification pipeline is not solely concerned with the significance of the dysconnected networks, rather whether or not information about chronotype is embedded within them such that classification can occur. Therefore, the threshold for significance was varied in the range [0, 1] in steps of 0.01 to understand its effect on the success of the classifier.

The effect of changing the significance threshold was considered for both the original datasets as well as the partial datasets, as mentioned in Section 2.9.

Note that a significance threshold of zero guarantees that every network is non-significant; therefore every classification is unclear due to step one. A non-zero but low significance threshold would result in the majority of subjects being classified due to step one - the significance of the network. As the significance threshold increases step two will be used for classification and finally once the significance threshold is high enough such that all 4 networks (*E_t_E__, E_t_L__, L_t_E__, L_t_L__*) are all significant then step three - the number of edges - is used. After reaching the significance level such that all four dysconnected networks are considered significant, the accuracy will remain constant for all significance levels higher than this. Therefore, the choice of significance threshold equates to the contribution each step of the classifier makes.

## 3 Results

Prior to applying the NBS method, we considered a group-level analysis comparing the graph metrics between the two extreme chronotypes was considered. No significant differences were found as shown in Appendix C, which is consistent with results found independently by Farahani et al (Farahani, Fafrowicz, Karwowski, Bohaterewicz, et al. 2021).

### 3.1 Classifier Performance

Having selected the t-statistic thresholds 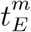 and 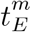 the classifier can be used as presented in Section 2.7. The results for each of the three scans under the two contrasts are presented below. In addition, the classifier labels assigned to each Subject are given in Table 3.

#### 3.1.1 Afternoon Scanning Session

For the contrast ECP>LCP, Subject 22 was mislabelled as an ECP on step one while all other subjects were labelled as unclear by step one, resulting in an accuracy of 0%. For the contrast ECP<LCP, accuracy was 0% with all subjects being classified by step one.

#### 3.1.2 Evening Scanning Session

For the contrast ECP<LCP, every subject was labelled as unclear by step one, due to 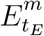 and 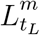 being non significant for all subjects, resulting in an accuracy of 0%.

For the contrast ECP>LCP, 97.37% accuracy was achieved due to only Subject 36 being misclassified as an ECP. The resulting subnetwork when all subjects are correctly labelled (i.e. NBS as presented in (Zalesky, Fornito, and Bullmore 2010) is used with t-statistic threshold 1.692) is given in Figure 4 and its topology is considered in Table 4. The breakdown of how many ECPs and LCPs were classified at each step is presented in Table 2.

#### 3.1.3 Morning Scanning Session

For the contrast ECP>LCP every subject was labelled as unclear by step one, due to 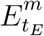 and 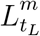 being non significant for all subjects, resulting in an accuracy of 0%. For the contrast ECP<LCP accuracy was 0%, with all subjects being classified by step one.

### 3.2 How the t-statistic Threshold Impacts Classifier Performance

As can be seen from Table 3, the performance of the classifier was strongly dependent on the time of day that the scan was taken, with accuracy differing greatly between the Evening and other scanning sessions. Since the accuracy of the classifier for the Afternoon and Morning scans was restricted by dysconnected networks being non-significant at step one of the classifier, when the percolation threshold was used, we now considered the accuracy of the classifier when selecting a single t-statistic threshold, varying over the range [0, 4.5] in increments of 0.01. This relaxes the requirement of connectedness when choosing the t-statistic threshold for defining the dysconnected network and allows an insight into the stability of the classifier when varying the t-statistic threshold. Under these conditions the classifier is the simplified version presented when 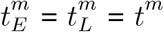 where *t^m^* is preselected.

First, we consider the accuracy values when applying the classifier for the three scanning sessions and two contrasts when varying the t-statistic threshold value. These results are presented in Figure 2. The highest accuracy values for the Afternoon scanning session occur for lower t-statistic thresholds near 2.2 when ECP<LCP (~ 97%). The contrast ECP>LCP does not result in any non-zero accuracy values. For the Evening scanning session, non-zero accuracy exclusively occurred for the ECP>LCP contrast. For this contrast, non-zero accuracy occurs between 0.71 and 2.28 with accuracy peaking at 97%. The Morning scanning session, non-zero accuracy was only seen for the ECP<LCP contrast, with t-statistic thresholds near 2.2 and 2.7 with accuracy ~ 90% for both thresholds.

**Figure 2:**
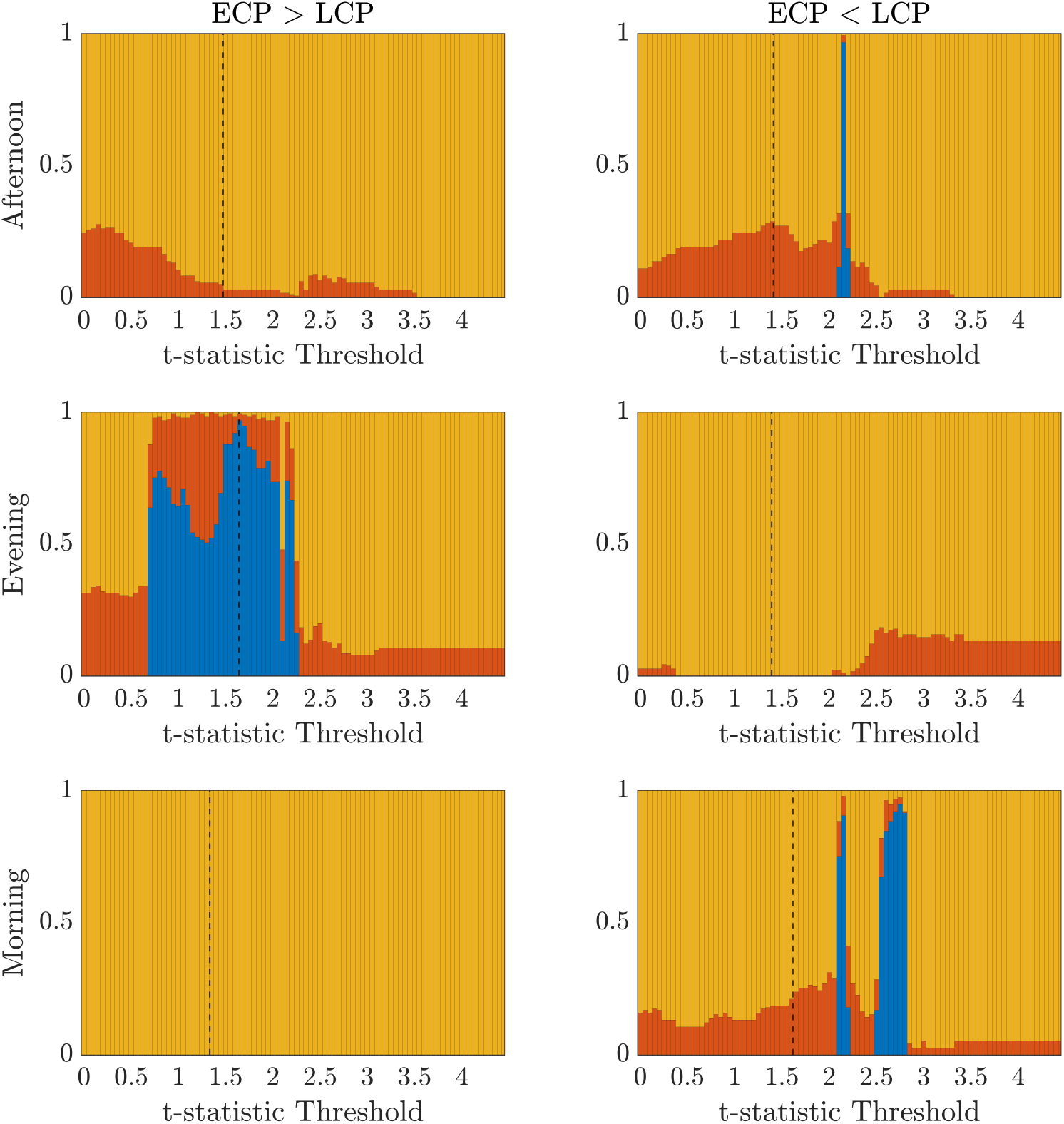
Classifier Performance as a Function of t-statistic Threshold. Stacked bar charts showing the percentage of subjects classified correctly, incorrectly and unclear respectively (blue, red, yellow) when successively selecting the t-statistic threshold from the range [0, 4.5] in increments of 0.01. The dashed lines indicate the percolation threshold for correct labelling as found using Section 2.6.

In Table 4 we display the topologies of four distinct dsyconnected networks corresponding to the regions of non-zero accuracy in Figure 2. In addition, Figures 3 - 6, visualise the four dsyconnected networks. For the Evening [ECP>LCP] scan the threshold of 1.692 is chosen as it is the percolation threshold when all subjects are correctly labelled as in Section 3.1.2. Since non-zero accuracy did not occur for the Afternoon [ECP>LCP], Evening [ECP<LCP] nor the Morning [ECP>LCP] scanning sessions, no topological features from these scans are considered. Due to the overlap in the t-statistic thresholds that produce high accuracy near at 2.2 in the Morning [ECP<LCP] and Afternoon [ECP<LCP] scanning session, these dysconnected networks at threshold 2.19 were compared for similarity. Notably, only one edge is present in both networks linking nodes 48 (LECN) and 64 (Visuospatial).

**Figure 3:**
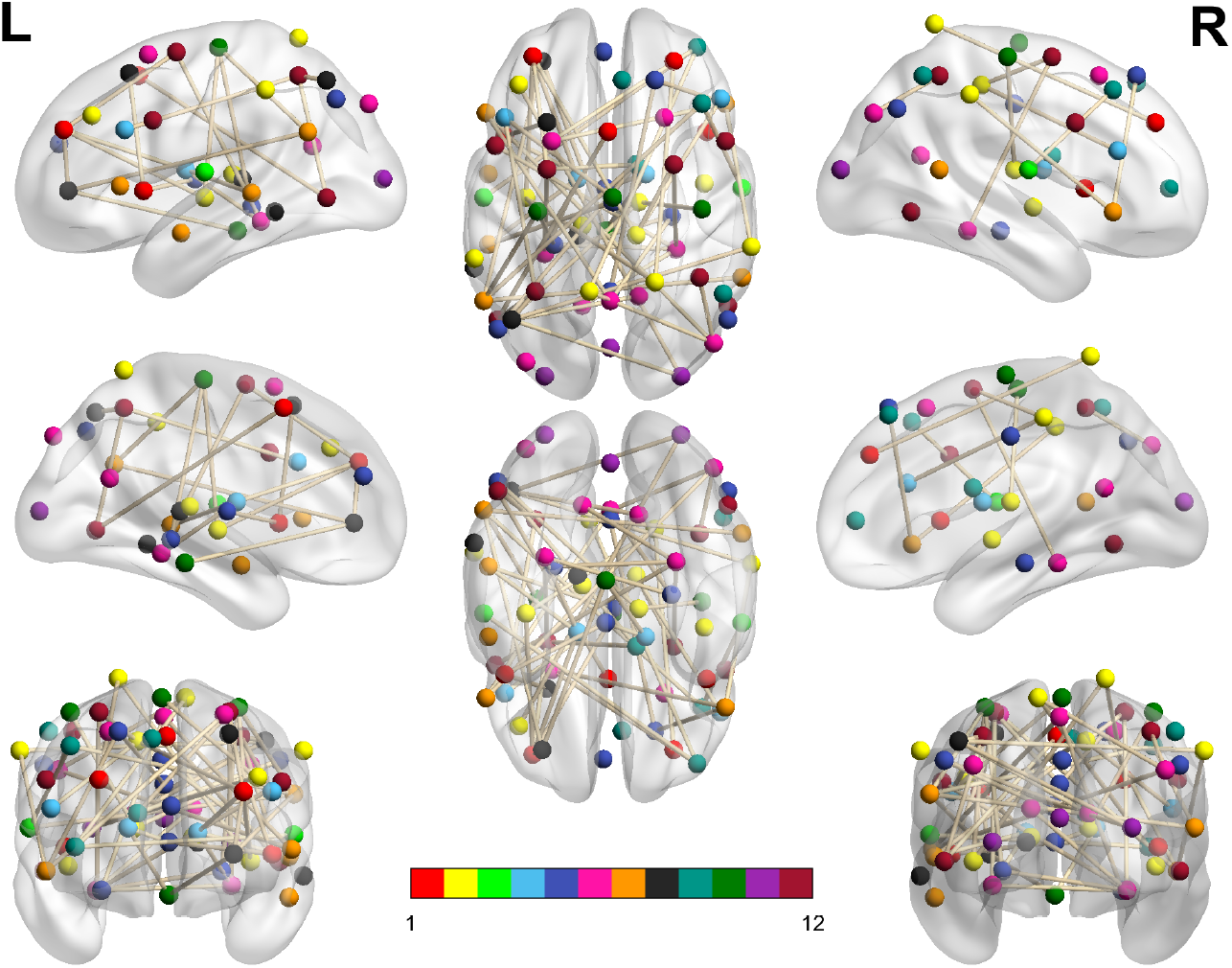
Dysconnected network produced using NBS for the Afternoon scanning session using ECP<LCP and t-statistic threshold 2.19. The top row from left to right are the lateral view of the left hemisphere, top view and the lateral view of the right hemisphere. The middle row from left to right are the medial view of the left hemisphere, bottom view and the medial view of the right hemisphere. The bottom row shows the frontal side on the left and the back side on the right. The interconnected networks (ICNs) 1-12 are given in Table 12.

**Figure 4:**
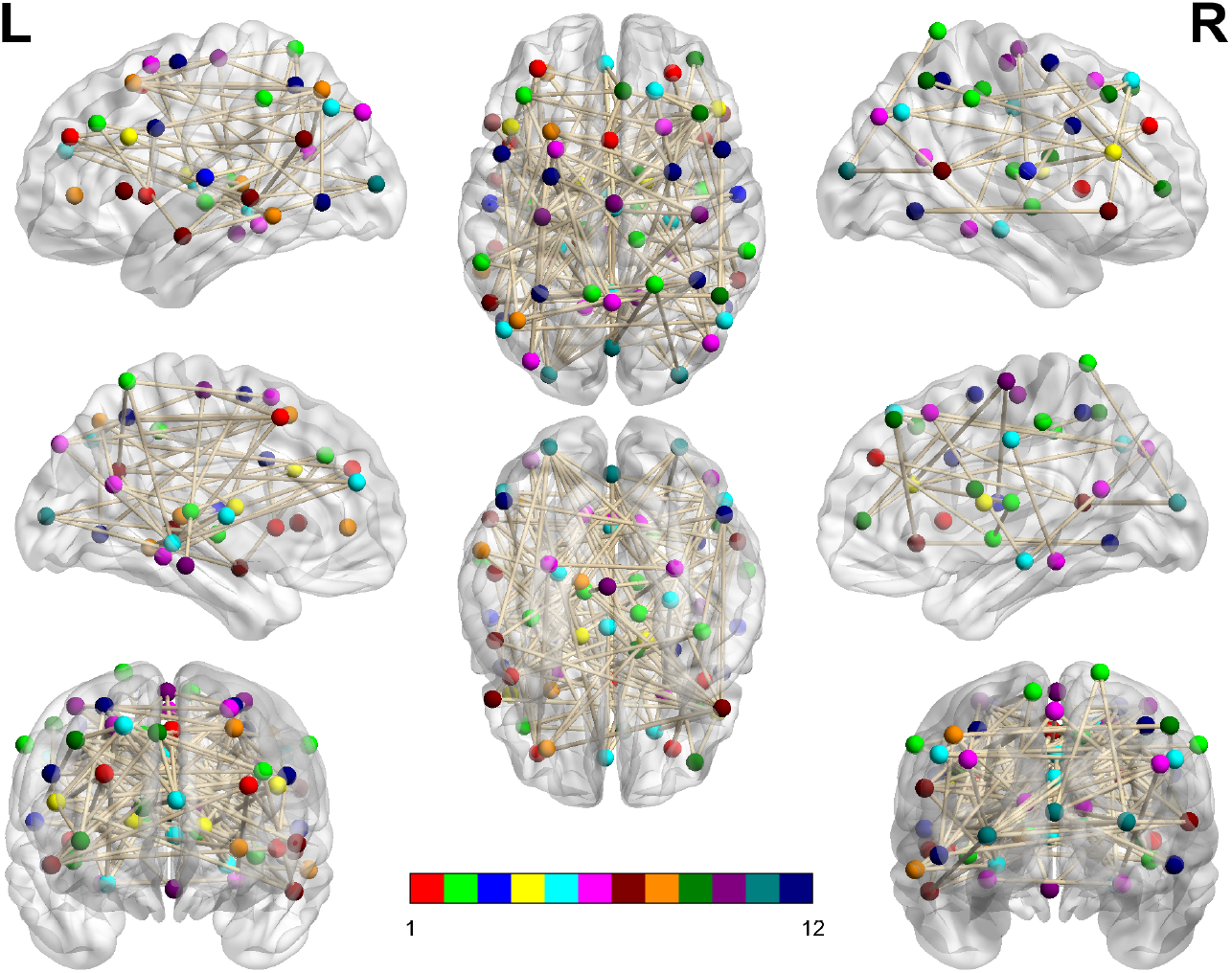
Dysconnected network produced using NBS for the Evening scanning session using ECP>LCP and t-statistic threshold 1.692. Subfigure information is the same as Figure 3.

**Figure 5:**
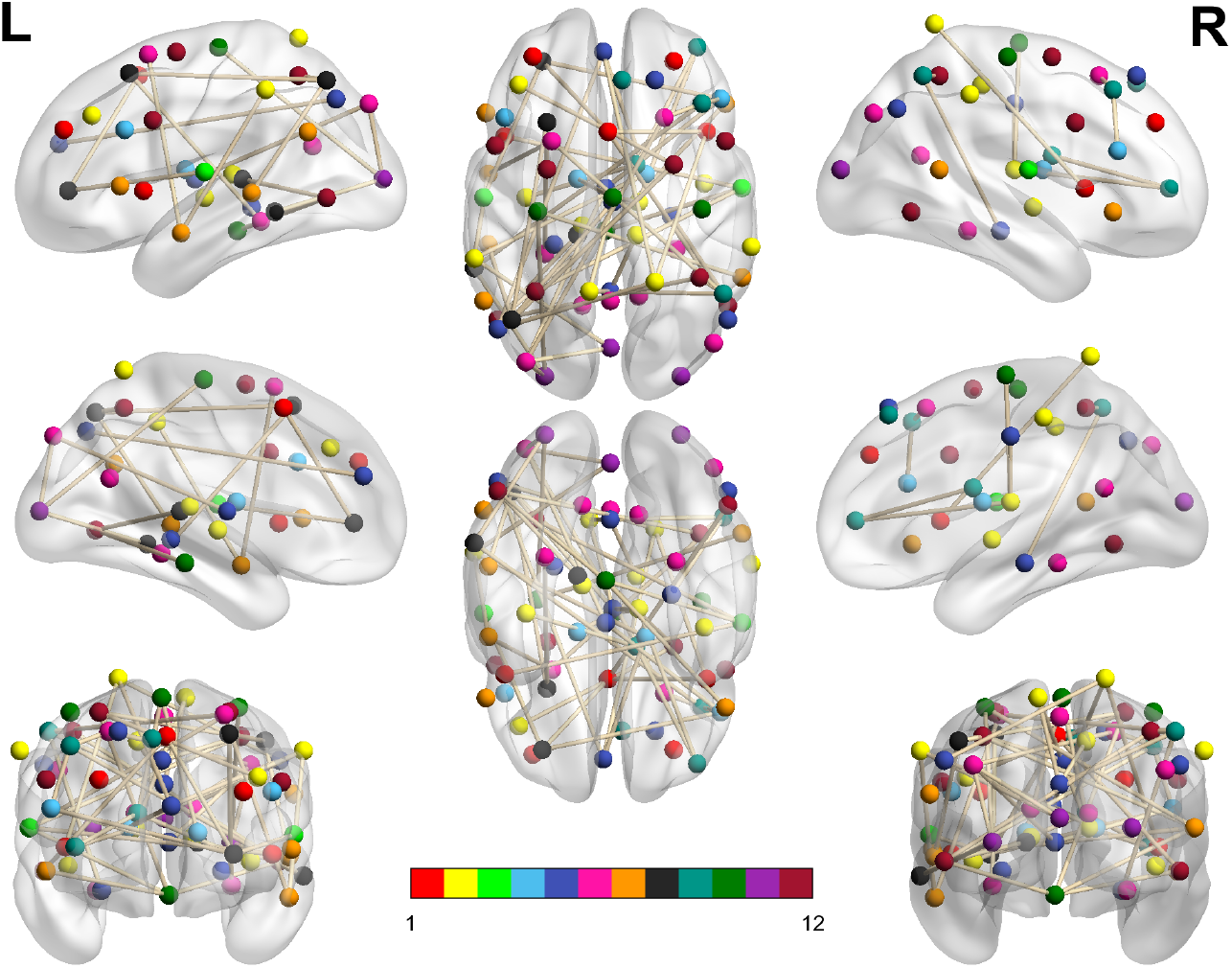
Dysconnected network produced using NBS for the Morning scanning session using ECP<LCP and t-statistic threshold 2.19. Subfigure information is the same as Figure 3.

**Figure 6:**
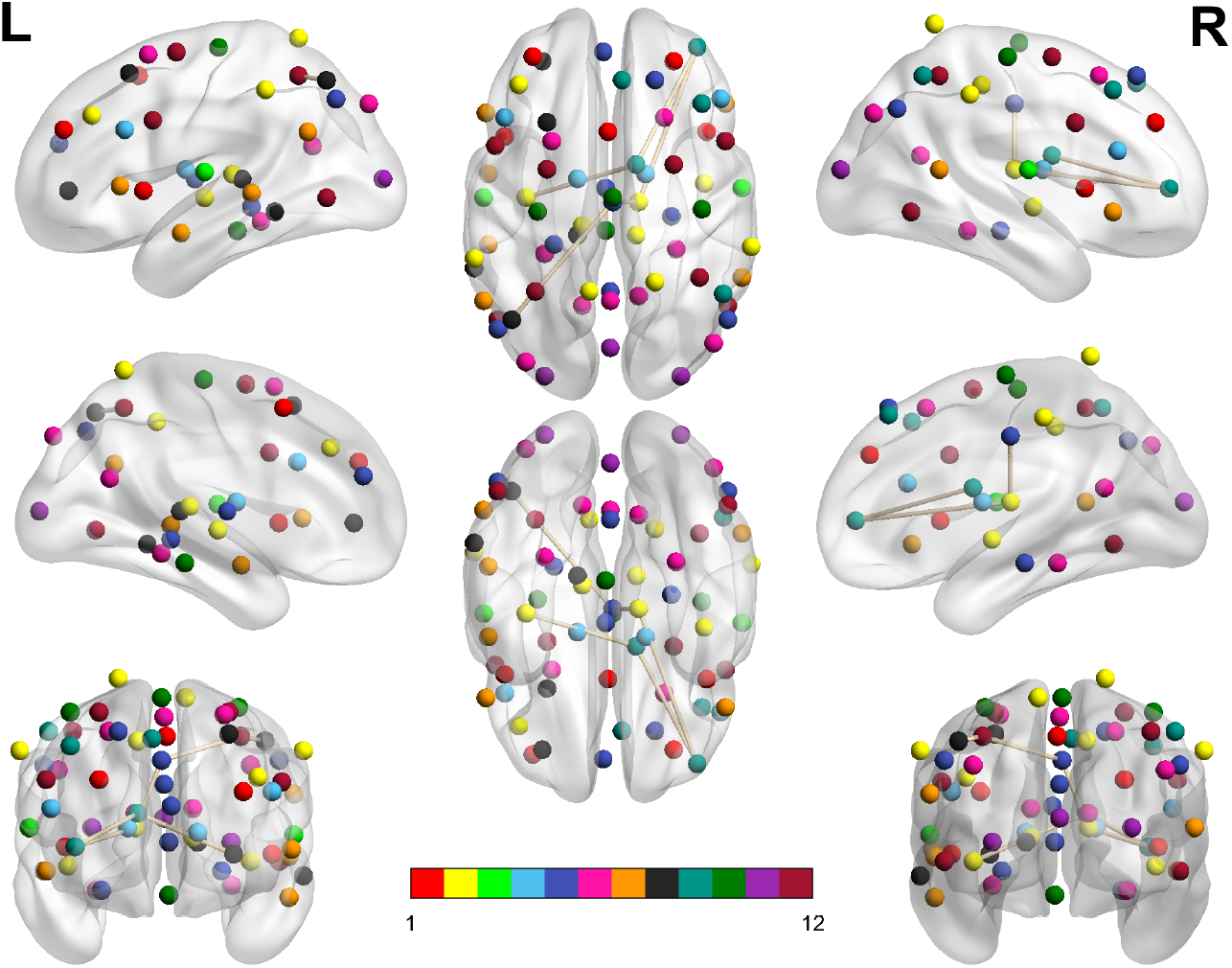
Dysconnected network produced using NBS for the Morning scanning session using ECP<LCP and t-statistic threshold 2.79. Subfigure information is the same as Figure 3.

It is important to note that the purpose of presenting these results is to provide insight into the somewhat unintuitive differences observed, dependent on time of day. We do not use this approach to construct a classifier, due to the issue of multiple comparisons introduced by varying the threshold over a wide range.

### 3.3 Investigating the Stability of the Classifier

So far the results of the classifier for the original datasets have been presented. However, to validate the classifier it is important to understand how the classifier performs on the partial datasets, created by sequentially leaving out participants. Therefore, the results of using the classifier, as presented in Section 2.9, on each of the new datasets created by removing one subject in turn, from both the training set and the test set, are presented below.

#### 3.3.1 Afternoon Scanning Session

The percentage of subjects who are labelled correct, incorrect and unclear when a particular subject is removed is shown in Table 5 and Table 6 for the contrasts [ECP>LCP] and [ECP<LCP] respectively.

For the contrast [ECP>LCP] accuracy was zero for all subjects due to a majority of subject classified as unclear. When comparing these results to the corresponding column in Table 3 we see a high consistency, as all but one subject was classified as unclear.

For the contrast [ECP<LCP] there are certain subjects (5, 6, 7, 10, 16, 23, 26 and 38) whose removal has positive effect on the accuracy of the classification. Indeed, except for Subject 38, the accuracy is higher than random chance (50%) and in some cases the classifier would be considered high performing. For example, the removal of Subject 16 sees accuracy go from 0% up to 81%.

When comparing the results to the corresponding column in Table 3 we see a high correspondence between the incorrectly labelled subjects in Table 3 and those subjects whose removal sees a high accuracy, especially for the ECP cohort.

#### 3.3.2 Evening Scanning Session

Table 7 and Table 8 show the percentage of subjects who were labelled correct, incorrect and unclear when a particular subject was removed, for the contrasts [ECP>LCP] and [ECP<LCP] respectively.

For the contrast [ECP<LCP] no subject was correctly labelled due to almost all subjects being classified as unclear. When comparing these results to the corresponding column in Table 3 we see extremely high consistency.

For the contrast [ECP>LCP] there are certain subjects (7, 8, 9, 16, 18, 19, 22, 25, 30, 32 and 34) whose removal results in a decrease in accuracy below random chance and far below the 97.3% seen for the full dataset. In addition, the mean accuracy of the other subjects is 79.68% - a reduction from the average of 97.3%. However, it is worth noting that the removal of Subject 36 - the only subject incorrectly classified in the original dataset - sees an increase in accuracy from 0% to 76%.

The low or zero accuracy for those eleven subjects is mainly due to the majority of the subjects being classified as unclear. There was no clear link to differences in the non-imaging data (e.g., actigraphy, DLMO etc) that could provide a reason for these subjects having such a clear influence on the classifier’s performance.

#### 3.3.3 Morning Scanning Session

The percentage of subjects who were labelled correct, incorrect and unclear when a particular subject was removed is shown in Table 9 and Table 10 for the contrasts [ECP>LCP] and [ECP<LCP] respectively.

For the contrast [ECP>LCP] all subjects were classified as unclear, which is consistent with the corresponding column in Table 3 where all subjects were classified as unclear.

For the contrast [ECP<LCP] there are certain subjects (3, 10, 13, 20, 26, 27, 29, 34, 36, 38) whose removal has a positive effect on the accuracy of the data. In the case of Subjects 3, 10, 13, 20, 26, 27 and 38 their removal produces accuracy’s higher than random chance (50%). For example, the removal of Subject 13 sees accuracy go from 0% up to 76%.

When comparing the results to the corresponding column in Table 3 we see a correspondence between the subjects whose removal sees a high accuracy and incorrectly labelled subjects in Table 3. However, Subjects 3 and 38 were not misclassified in Table 3 whilst Subjects 2, 19, 36 and 37 were misclassified in Section 3.1.3 but their removal had no discernible effect on the classifier.

### 3.4 How the Significance Threshold Impacts Classifier Performance: Partial Datasets

Due to the sensitivity of the classifier, both in terms of removing subjects resulting in increased accuracy in the Morning and Afternoon scanning session [ECP<LCP] and the reduction in accuracy in the Evening scanning session [ECP>LCP], the influence of the choice of significance threshold was investigated.

It is worth noting that the initial investigation into the classifier’s sensitivity to the removal of subjects focused on the subjects themselves. Therefore, an investigation into the metadata of subjects who were misclassified or whose removal led to high differences in accuracy was undertaken. However, as seen in (Appendix D) the metadata is unable to provide an answer for the classifiers’ sensitivity.

Figure 7 shows boxplots for all of the n accuracies when one subject in turn has been removed from the training and test set for all possible values of the significance threshold *α*. In addition, the mean and standard deviation for each *α* value are shown. Figure 8 shows the mean value for the percentage of subjects labelled correct, incorrect and unclear for each *α* value.

**Figure 7:**
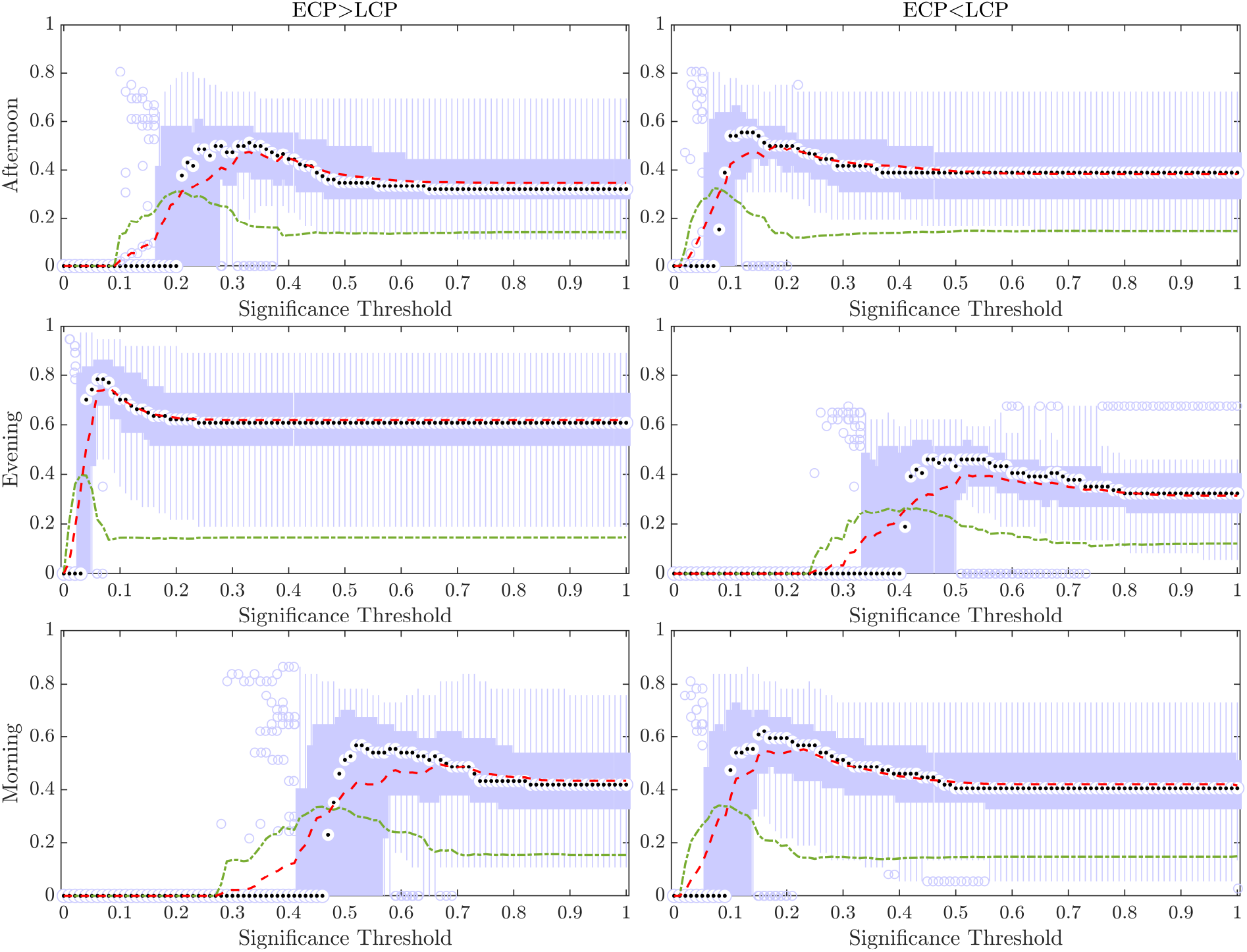
Boxplots Showing the Accuracy When Removing Subjects and Varying Across Thresholds for Significance. For each value of significance, in the range 0 to 1 in increments of 0.01, a boxplot for the accuracy when removing each subject is shown as as the mean accuracy (red, dashed) and the standard deviation (green, dashed and dotted). The median of the boxplot is given by the black dot while values considered outliers (greater than 2.7 standard deviations away from the mean) are depicted by small blue circles.

**Figure 8:**
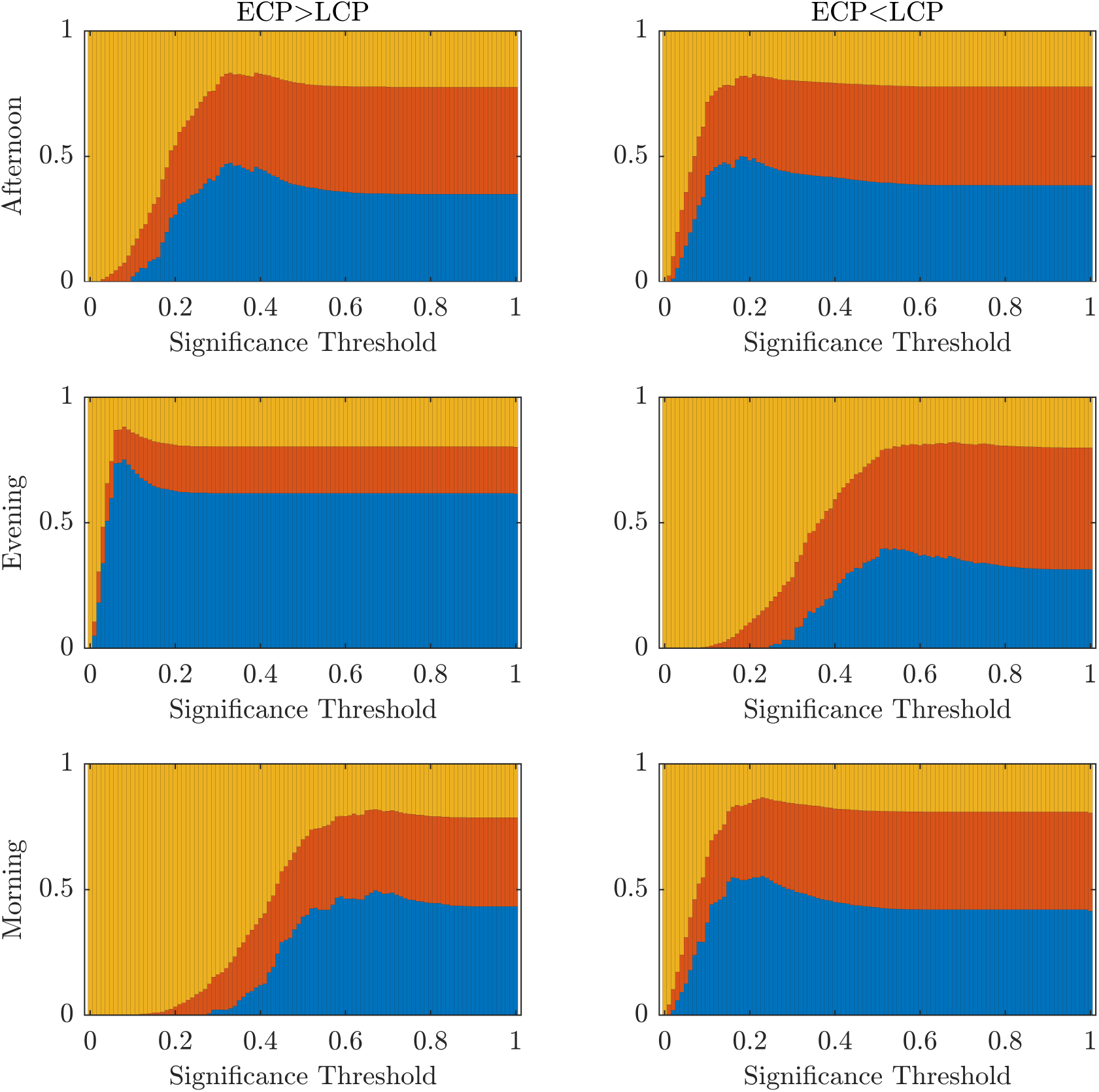
Performance of the Classifier When Removing Subjects and Varying Across Thresholds for Significance. Stacked bar chart showing the mean percentage of subjects classified correctly, incorrectly, and unclear respectively (blue, red, yellow) when removing one subject in turn and varying across the threshold for significance.

As can be seen from the Figures 7 there is an optimum value of *α* for each of the scanning sessions, selected as the point where the ratio between accuracy and standard deviation is highest.

However, it is only in the Evening [ECP>LCP] scan that the value of *α* which results in the maximum accuracy matches the value of *α* which produces the minimum standard deviation. This occurs at *α* = 0.08 resulting in a mean accuracy of 75.25% showing that there is the possibility to optimise the significance threshold for *α* and that selecting the traditional significance level of *α* = 0.05 may be too conservative when there are additional steps of the pipeline that help to prevent misclassifications.

### 3.5 How the Significance Threshold Impacts Classifier Performance: Original Datasets

To understand how changing the significance threshold effects the performance of the classifier when considering the original datasets, the results when varying the significance between [0, 1] in increments of 0.01 are shown below.

Figure 9 shows a stacked barchart showing the distribution of correct, incorrect and unclear classification of each possible value of *α* when ranging from 0 to 1 in increments of 0.01 for the original full datasets. The maximum accuracy of 97% occurs for the Evening [ECP>LCP] scan for *α* in the range 0.04-0.07. Comparing Figure 9 to Figure 8 we see similar trends with accuracy increasing with increasing *α* up to a point specific to each scan and then steadily declining to a plateau once *α* is high enough. Table 11 shows the values of *α* for which the maximum accuracy occurs.

**Figure 9:**
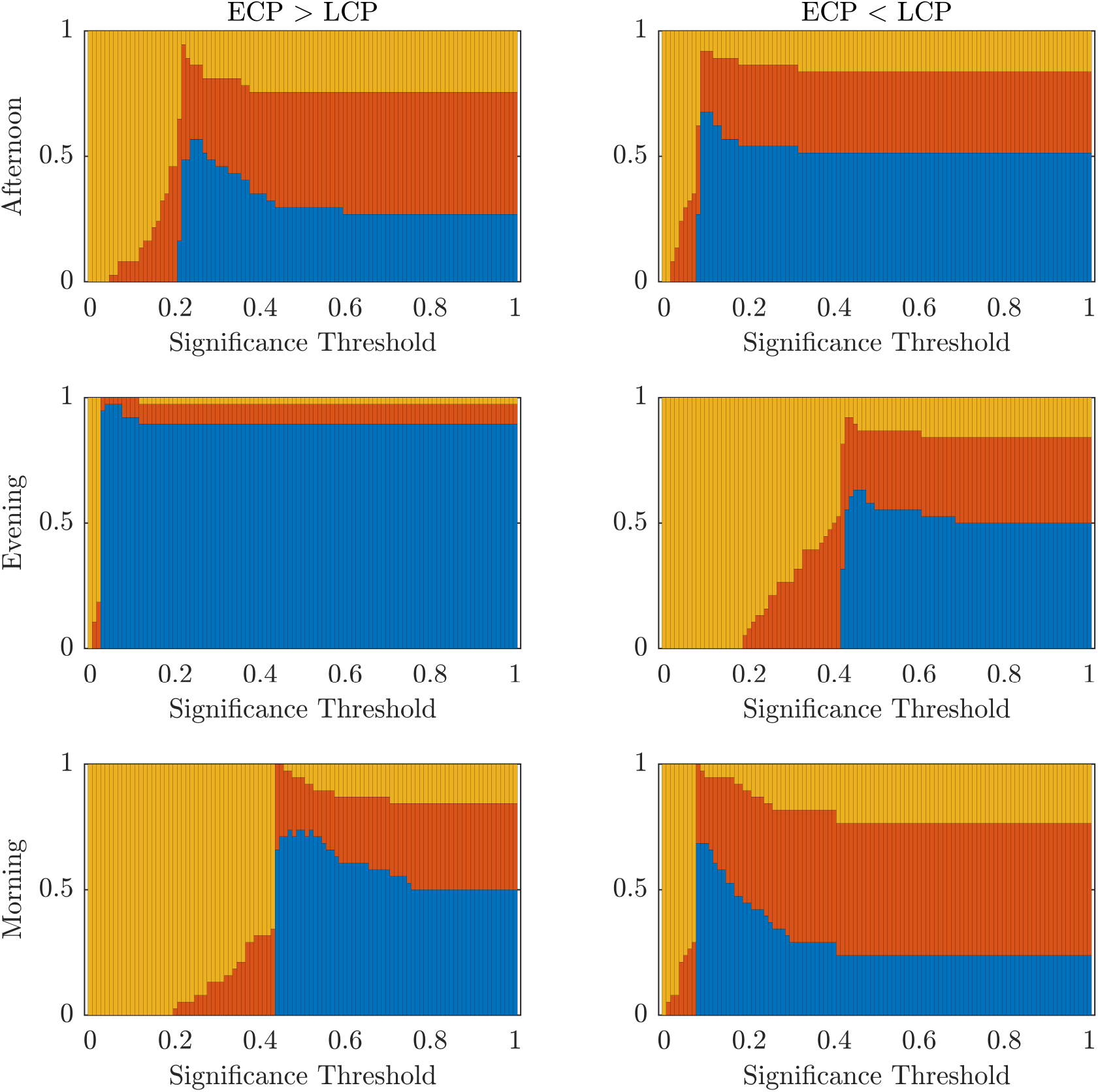
Performance of the Classifier When Varying Across Thresholds for Significance. The percentage of subjects classified correctly, incorrectly, and unclear respectively (blue, red, yellow) when varying across the threshold for significance.

## 4 Discussion

To the best of our knowledge, this is the first study explicitly aiming to classify an individual’s chronotype using rs-fMRI data. While previous studies (Elise R. Facer-Childs, Brunno M. de Campos, et al. 2021; Elise R. Facer-Childs, Brunno M. Campos, et al. 2019; C. M. Horne and Norbury 2018) have identified differences in functional connectivity associated with chronotype, the question of whether these differences are sufficient to identify a participant’s chronotype solely from fMRI data has not been asked. Following creation of FNs, NBS is used as the base for a classifier. The classifier is innovation through its evaluation of whether an ECP or LCP classification of the test subject leads to a clearer differentiation between the two classes in a group-level comparison.

In addition, the classifier was presented alongside a principled way to select the t-statistic thresholds, a criticism of NBS. This focused on the two percolation thresholds resulting from the two different chronotype labellings a test subject can be assigned. Through concentrating on edges located in dysconnected subnetworks there is evidence that rs-fMRI data does contain enough information to distinguish between ECPs and LCPs. Indeed, for the Evening [ECP>LCP] scan we see high classification accuracy of 97.3% when applying the percolation thresholds.

The high level of accuracy is predominantly due to step one of the classifier, which compares the significance of the MCCs 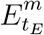 and 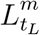. This step contributed almost one third of the correct classifications, as seen in Table 2. Since the MCC, when all subjects are correctly labelled, covers all 70 ROIs with only 143 edges (Table 4), it suggests that the key differences between the FNs of extreme chronotype are sparse and distributed during the evening.

This may explain why neither the seed-based approach in (Fafrowicz et al. 2019), nor the graph metric approach in (Farahani, Fafrowicz, Karwowski, Bohaterewicz, et al. 2021) could find significant differences between extreme chronotypes. Since the effect of chronotype is subtle and seen only through the group level comparison of specific edges using a contrast to compare between ECPs and LCPs, this would be lost in many graph metric approaches, which average over many nodes or the entire network reducing the focus on key edges. On the other hand, seed-based approaches that rely on contrasts may identify differences (Elise R. Facer-Childs, Brunno M. de Campos, et al. 2021; Elise R. Facer-Childs, Brunno M. Campos, et al. 2019), but the focus on specific seeds could limit the ability to identify the distributed effect seen in the Evening [ECP>LCP] scan.

Meanwhile, the five results of 0% accuracy for the chosen thresholds raises questions about why the classifier’s performance is so optimised for the Evening scan and why scans at other time points seem to have no ability to differentiate between ECPs and LCPs. While some variation in accuracy throughout the day may be expected, especially since time of day effects have been linked to chronotype (Fafrowicz et al. 2019), such contrasting results have no apparent explanation in chronobiological literature.

A key assumption in the pipeline is that all ROIs provide valuable information and the effect of chronotype is therefore distributed throughout the brain. However, due to the poor performance across the different scanning session this assumption seems unsuitable for the Morning and Afternoon scans. Previous studies using the same data (Elise R. Facer-Childs, Brunno M. Campos, et al. 2019; Elise R. Facer-Childs, Brunno M. de Campos, et al. 2021) have suggested that the differences in FC associated with chronotype and time of day can be restricted to individual nodes, or even sub-portions of individual nodes. The optimal way to define a node (and related issues such as the optimal number of nodes to use etc.) therefore remains an issue and its impact on the classifier requires further research (Korhonen, Saarimäki, et al. 2017; Korhonen, Zanin, and Papo 2021; Song, Panych, and Chen 2016).

The choice of the t-statistic threshold was investigated to understand the restriction of the MCC assumption. Figure 2 shows it is possible at all three times of the day - under one contrast - to find a t-statistic threshold that results in a highly accurate classifier for distinguishing between ECPs and LCPs. This expands the results presented in Section 3.1, which focus on a specific threshold. However, we restricted our analysis to cases where a principled approach could be taken to defining the threshold, rather than highlighting situations where good classification accuracy could be achieved with an arbitrary threshold. Clearly, the high classification accuracy in these cases also supports the view that chronotype classification can be undertaken with rs-fMRI data, although further work is needed to develop the statistical methods to achieve this without the use of arbitrary thresholds.

Despite the problem of statistical significance arising from multiple comparisons, Figure 2 does offer some insights into the effect of chronotype over the day and especially in relation to the classifier. For instance, the contrast resulting in high accuracy, when varying the t-statistic threshold, changes over the course of the day. Simplistically, this generally follows the pattern of high classification accuracy occurring when the chronotype with increased tiredness (as measured using KSS (Elise R. Facer-Childs, Brunno M. Campos, et al. 2019)), has a higher FC in the contrast. Hence, the contrast ECP>LCP performs better in the evening when ECPs are likely to be more tired than LCPs. Similar logic follows for the Morning scan, when forcing LCPs to awaken before their natural sleep pattern for an 08: 00 scanning session will result in increased tiredness for that cohort. Finally, the contrast ECP<LCP produces non-zero accuracy in the Afternoon. However, the range of t-statistic thresholds, which produce non-zero accuracy, is smaller in the Afternoon compared to the Morning. This could be associated to the greater similarity in KSS scores in the two groups at this time. This may suggest that the classification is driven by tiredness (or Process S within the two process model (Borbély and Achermann 1999)) rather than chronotype (related to Process C) per se. Within a real-world setting, and not only in relation to rs-fMRI data, differentiating the impacts of sleep homeostasis and circadian drive is difficult. Future studies using laboratory based constant routine or forced desynchrony protocols (Duffy and Dijk 2002; Kyriacou and Hastings 2010) could help to understand how sleep drive and circadian phase are differentially manifested in rs-fMRI data and brain networks more broadly.

Furthermore, the t-statistic thresholds producing high accuracy could provide an insight into how chronotype affects the brain throughout the day. Indeed the size of the dysconnected networks producing high accuracy is markedly different throughout the day. The large range of t-statistic thresholds in the Evening start at 0.71, representing a dense distributed effect, while the highest peak in accuracy near 1.7 suggests a sparse but distributed effect. In contrast, in the Afternoon and Morning the high t-statistic thresholds near 2.2 suggest a focal effect concentrated on specific subsets of the brain’s FN. Finally, the second peak in the Morning near 2.8 is focused on only 7 nodes and therefore represents a highly spatially specific impact on the brain. More detailed investigation of the spatial distribution of network changes as a function of time of day and chronotype would help to develop these ideas and provide a more specific understanding of how the brain is impacted.

The change in the range of t-statistic thresholds that result in high accuracy explains why the classifier only saw results of 97.3% for Evening [ECP>LCP] scan, and 0% for the other scans and contrasts. This is because it is the only combination where the percolation threshold when all subjects are correctly labelled, shown by the dashed line in Figure 2, falls directly within the range of t-statistic thresholds that consistently sees non-zero accuracy. For the other five combinations this is not the case. This indicates that the assumption of connectedness, the focus on MCCs and the conventional choice of *α* = 0.05, is suitable for the Evening [ECP>LCP] scan, while a different combination of parameters needs to be used to optimize accuracy for the other scans. Given our results and the discussion above, other approaches could be developed to extract the information needed to provide good classification accuracy in the Morning and Afternoon. Our work would suggest that the information is present in the data.

In addition, the stability of the classifier was investigated, through the creation of additional surrogate datasets using a leave-one-out approach. It is clear from the results across all 3 scanning session, as seen in Tables 5 - 10, that the classifier is highly sensitive to the removal or inclusion of certain subjects. In the case of the Afternoon and Morning scanning sessions the removal of certain subjects led to a considerable increase in accuracy, while in the Evening scanning session subject removal has the opposite effect, reducing accuracy, aside from Subject 36.

We see that when comparing the results from the partial datasets to the original datasets there is a correspondence between improved accuracy when incorrectly labelled subjects in Table 3 are removed. This may indicate that these subjects have properties closer to the other chronotype, and hence why their removal improves the classifier. Similarly, we might hypothesise that the near zero accuracy resulting from the removal of subjects in the Evening scanning session could be because they have properties which clearly identify them as their phenotype and hence their removal reduces the accuracy of the classifier. However, this hypothesis was not supported by differences in the metadata as shown in Appendix D. Similarly, the metadata offers no insight into why, for example, Subject 36 was misclassified in the Evening [ECP>LCP] scan. Further work is needed to understand the subtleties of the links between resting brain function measured with fMRI and more established markers of chronotype.

Furthermore, it was observed that a large proportion of the subjects were classified as unclear due to non-significant networks occurring at the traditional 0.05 significance level when the number of subjects was reduced by one. To investigate if differentiable information is present when using higher significance thresholds an investigation into the significance threshold was also completed.

The accuracy of the classifier as well as the standard deviation of the *n* different dataset’s accuracy for each significance threshold are presented in Figure 7. An optimal choice of significance threshold could be selected for each scanning session and contrast to maximise the accuracy across the *n* datasets and minimise the standard deviation. This relationship is the clearest for the Evening scanning session and indeed using *α* = 0.08 shows the improved ability to differentiate chronotypes in the Evening scanning session. For the other scanning sessions and contrasts this relationship is not as distinct, but an optimal ratio between these two factors could be located. Directly relating the parameter choice for significance to the classifier’s stability offers a solution for how to improve the classifier’s performance in future datasets.

When comparing the optimal significance thresholds seen in Figure 7 for the partial datasets, to the optimal values in Table 11 for the original datasets, we clearly see that *α* = 0.05 is too conservative for *n* – 1 subjects compared to *n* subjects. One possible explanation for this is that the number of subjects is quite low and reducing the size further removes important information. Indeed, the reduction in the optimum value for α decreases for the Evening [ECP>LCP] for 38 subjects (0.04-0.07) compared to when 1 subject has been removed (0.08), indicating that as the number of subjects increases, the stability of the classifier increases. However, it is reasonable to assume this trend will not continue indefinitely and that there will be an optimum number of subjects in the training set such that they provide enough information for classification, while also allowing the test subject to have enough influence that changing their label will have an effect on the t-statistic. This highlights that the classifier is reliant on the fact that changing the label of one subject will lead to a detectable impact on the t-statistics, while also having a large enough training set to ensure there is enough distinction between the two cohorts. This approach is optimised to smaller datasets, where mislabelling one subject will have a greater influence. If the number of subjects increased sufficiently (n → ∞) it is reasonable to assume the dysconnected networks produced will be the same irrespective of the labelling of the test subject. At that stage this classifier would be redundant and NBS could be used in its traditional form.

For both the partial datasets in Figure 8 and the original datasets in Figure 9, we see the optimal values of significance threshold follow a trend similar to that observed in Figure 2. Indeed, we clearly see high differences in the optimal value of α for the different contrasts in the Evening and Morning scan with the optimal significance threshold range being lower when the contrast aligns with the group who are experiencing increased tiredness as measured using KSS (Elise R. Facer-Childs, Brunno M. Campos, et al. 2019). In the Afternoon, while there is a difference in the optimal parameter choice between the two contrasts in agreement with KSS, the difference between the optimum α value in the two contrasts is smaller, which again could be associated to both groups having a more similar state of tiredness.

This pattern matches studies where excitability as measured through transcranial magnetic stimulation is higher when participants are more tired (Huber et al. 2012; Ly et al. 2016). In addition, (Petkov et al. 2014) showed that the propensity of people with epilepsy to transition into a seizure state is greater for networks with higher mean degree, which are therefore considered more excitable. If we view higher FC as a proxy for larger mean degree then the pattern of scanning session, contrast and accuracy would be linked to excitability and consequently tiredness of the two chronotype groups. This suggestions remains to be investigated in more detail, potentially with studies involving explicit sleep deprivation.

Overall, optimising the parameters of the classifier, which include two thresholding choices, should be the aim of future research. Both the value at which to threshold the t-statistic matrix and the threshold for significance must be selected. In both cases the selection of this value is somewhat arbitrary, and typically determined by convention. By varying over these thresholds it becomes clear that there is differentiable and important information present in the Afternoon and Morning scanning sessions and that optimising these parameters will improve the accuracy and importantly the sensitivity of the classifier to new datasets. Further research is needed to understand how a rigorous selection process or different underlying assumption could result in selecting thresholds that optimise accuracy for the Morning or Afternoon scanning session. Such research may also develop new ways in which fMRI can be used in the study of chronotype as well as lead to greater insight into the impact of chronotypes on brain FNs. This study also motivates the future use of other methods for quantifying brain function to investigate human chronotype. For instance, assessing the pipelines’ suitability for use with EEG data would be a natural extension, especially since MCCs were originally shown to be useful at detecting subtle effects on FNs from EEG recordings (Vijayalakshmi et al. 2015). This may result in the Morning and Afternoon scans having non-zero accuracy for the percolation thresholds, offer other interesting insights or simply increase the practically of recording sessions in relation to cost and location. However, compared to fMRI, EEG has limited spatial sampling of the brain, and a lack of sensitivity to deep brain structures which are known to be important to sleep and circadian regulation.

Furthermore, this study allowed subjects to sleep using their preferred schedule for two weeks prior to the scans. It is currently unknown if the differentiation and accuracy seen at the three scanning times would stay the same if LCPs were constrained to a more traditional work schedule or if high accuracy would occur at a different time of the day. It seems likely that with the additional sleep pressure associated with conforming to the societal day, LCPs would be more easily differentiable from ECPs.

## 5 Conclusion

In conclusion, we have shown that extreme early and late chronotypes have differentiable information in their rs-fMRI data that can be used to classify them. Indeed, this classifier demonstrates that two groups of participants whose differences are relatively subtle (i.e., not based on a clinical diagnosis) can be differentiated using rs-fMRI, when traditional seed-based and graphical methods have struggled.

Through this study we have proposed a classifier and investigated its sensitivity and robustness to changes in parameters and the training set. In an ideal scenario the removal of individual subjects would have a negligible effect on the classification accuracy. However, we found the accuracy of the classifier to be strongly dependent on individual participants, although these participants did not appear to be unusual based on the physiological and behavioural data we had available. We found conditions under which this could be mitigated by altering a combination of the threshold for the t-statistic, significance threshold for the dysconnected network and contrast being used. For future chronotype datasets it is recommended to use a contrast which reflects the tiredness of the two groups at the time of the scan, while issues around the optimal thresholds for significance and t-statistic thresholds remain to be clarified.

## Acknowledgments

Sophie L. Mason acknowledges the financial support of the University of Birmingham through an Alumni Scholarship. Leandro Junges acknowledges support from a Waterloo Foundation research grant (Ref no. 1970-4687). Wessel Woldman acknowledges the financial support of Epilepsy Research UK through an Emerging Leader Fellowship (F2002). Brunno M. Campos acknowledges the financial support of Sao Paulo Research Foundation (2013/07559-3). Elise R. Facer-Childs acknowledges financial support for this work from the Biotechnology and Biological Sciences Research Council (BBSRC, BB/J014532/1). E.R.F-C has received funding from the Department of Industry, Innovation and Science, Australia, #ICG000899 and #ICG001546) and is currently supported by a Science Industry Endowment Fund Ross Metcalf STEM+ Business Fellowship administered by the Commonwealth Scientific and Industrial Research Organisation. E.R.F-C and Andrew P. Bagshaw acknowledge the Engineering and Physical Sciences Research Council (EPSRC, EP/J002909/1) and an Institutional Strategic Support Fund Accelerator Fellowship from the Wellcome Trust (Wellcome, 204846/Z/16/Z). John R. Terry acknowledges the financial support of UKRI (EPSRC) via Fellowship EP/T027703/1.

## Data Availability Statement

De-identified participant data can be made available by contacting the corresponding author upon request. Reuse is only permitted following written agreement from the corresponding author and Primary Institution. The code used to create Tikhonov Partial Correlation Matrices as well as the classifier is available online at the github repository https://github.com/sophie-l-mason/NBS_classifier_code.

## A ROI Information

## B Derivation of Tikhonov Partial Correlation Matrix

Calculation of Tikhonov partial correlation is given below, extended from an outline provided in Pervaiz et al. 2020.

Let 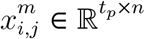 be the *i*,*j*^th^ component of data matrix *X*^(*m*)^ for Subject *m* for a rs-fMRI scan with *t_p_* time points and *n* ROIs. Then for the *m*^th^ participant an emperical covariance matrix, 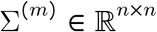, can be calculated using,

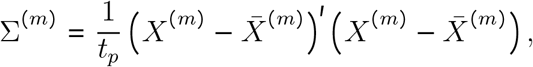

where ^′^ denotes matrix transpose and 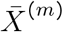 is the matrix created from vertically stacking the temporal average for each ROI *t_p_* times,

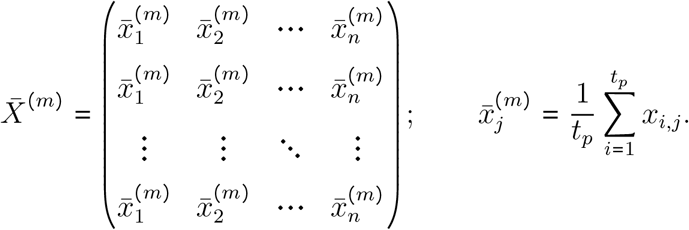

Now the Tikhonov covariance matrix is given by,

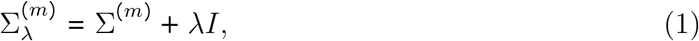

where 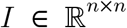 is an identity matrix. The positive constant λ controls the strength of the regularisation while the matrix *I* acts as the target matrix during the inversion process (Kuismin and Sillanpää 2017). Therefore, λ = 0 will result in partial correlation whilst higher values of λ will increasingly force off-diagonal elements closer to 0 and diagonal elements closer to 1. The Tikhonov precision matrix, 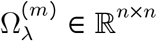, with elements 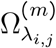 or to simply notation 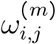, can then be calculated as,

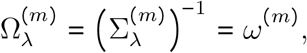

using Cholesky decomposition as explained in (Krishnamoorthy and Menon 2013) to computationally calculate the inverse of the Tikhonov covariance matrix using code provided by E. Blake (Blake 2015). Finally, the Tikhonov partial correlation matrix (TPCM), can be calculated element wise for *i* ≠ *j* such that each element is given by,

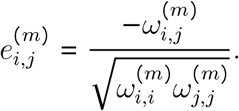

Partial correlation by definition requires two variables to compare whilst controlling for at least one other, therefore *i* = *j* and the resulting *e_i,i_* = −1 is meaningless. Hence, values on the main diagonal of the TPCM have been set to NaN and no self-links can be identified by this method.

The values in the TPCM are the FC between different brain regions and can be used as weights for the strength of statistical connections between different ROIs. If *e_i,j_* = 0 then under the assumption of fMRI data having Gaussian noise (Heras and Margalef 2013) there will be no edge between ROIs *i* and *j*.

The selection of the optimal regularisation parameter λ in eq(1) is based upon minimising an objective function.

First, let 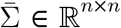 be the average empirical covariance for all N participants calculated by taking the element wise mean across Σ^(*m*)^ for the *N* participants. Also, let 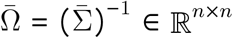 be the precision matrix associated with the average empirical covariance matrix, with elements 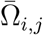. A matrix 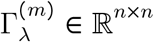 with elements, 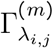, can then be defined using,

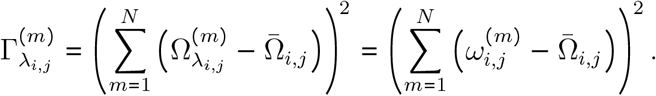

Each element is calculated by summing the difference between the *i*, *j*^th^ element of the average precision matrix and each participant’s corresponding λ specified Tikhonov precision matrix element before squaring the result.

Therefore, the optimal value of λ denoted λ_opt_ is the one which minimises the square root of the sum of the upper triangular elements in Γ_λ_ and hence is selected for Tikhonov partial correlation,

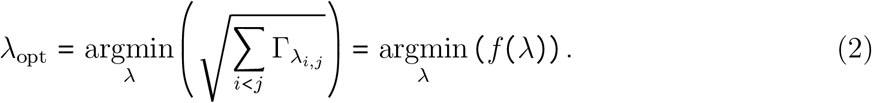

Potential λ values in the range [0.0001,1] in steps of 0.0001 are considered. However, for new datasets it would still be important to check the data is standardised, λ_opt_ is unique and that λ_opt_ ≠ λ^upper^. The value of λ^upper^ = 1 has been artificially set based on the data available however if λ_opt_ = λ^upper^ it would be important to understand the behaviour of *f*(λ), to ensure that the argmin has actually been found. A simple example that highlights the importance of this is that if *f*(λ) is a monotonically decreasing function then a search range of [0, λ^upper^] would always result in λ_opt_ = λ^upper^. Increasing λ^upper^ would therefore see λ_opt_ change accordingly despite no optimum actually existing.

Note that this provides one value of λ for which to regularise the *N* participants being considered. In this case one value of λ was optimised across all 113 scanning session across the 3 scanning times.

## C Traditional Graph Metric Approach

benInitially, group level analysis was considered using a traditional graph metric approach. The FNs, as constructed in Section 2.4 are used for the base of this analysis.

### C.1 Thresholding and Binarizing

Within this study the FNs are thresholded and binarized, such that values below the threshold are set to 0 and values above the threshold are set to 1. The threshold for binarizing the edges ranged incrementally in steps of 0.01 between the lower limit of 0 and the upper limit given by the percolation threshold. The minimum percolation threshold across subjects was selected as the maximum threshold to ensure all networks were connected at all thresholds considered. This creates more physiologically accurate networks, as different brain regions are known to be connected both structurally and functionally.

### C.2 Graph Metrics

After the networks have been thresholded and binarized graph metrics were calculated for each network. A limited number of graph metrics ranging across local and global measures were used. These are node degree, node strength, betweenness centrality, clustering, local efficiency, global efficiency, characteristic path length, assortativity and small-world index and small-world propensity. The graph metrics were calculated using using the freely available Brain Connectivity Toolbox (Rubinov and Sporns 2010) as well as (Humphries and Gurney 2008) for small-worldness. In the case of local measures the mean across all nodes was considered. It is worth noting these graph metrics align very closely to those used in a concurrent chronotype study completed by (Farahani, Fafrowicz, Karwowski, Bohaterewicz, et al. 2021.

### C.3 Group-Level Analysis

Finally, a group level analysis was completed by considering if the graph metrics for the two groups are significantly different at a given threshold value. Significance testing between the two groups distributions were completed using permutation testing (*n* = 10, 000 permutations). In addition, corrections for multiple comparisons across graph metrics as well as thresholds were completed using both a Bonferroni correction and the less conservative Benjamini-Hochberg correction.

### C.4 Results

The distributions for the different graph metrics for each group are given in Figures 10 to 12 for the three scanning sessions when thresholding and binarizing the whole brain network in increments of 0.01 from 0 up to their MCC threshold.

**Figure 10:**
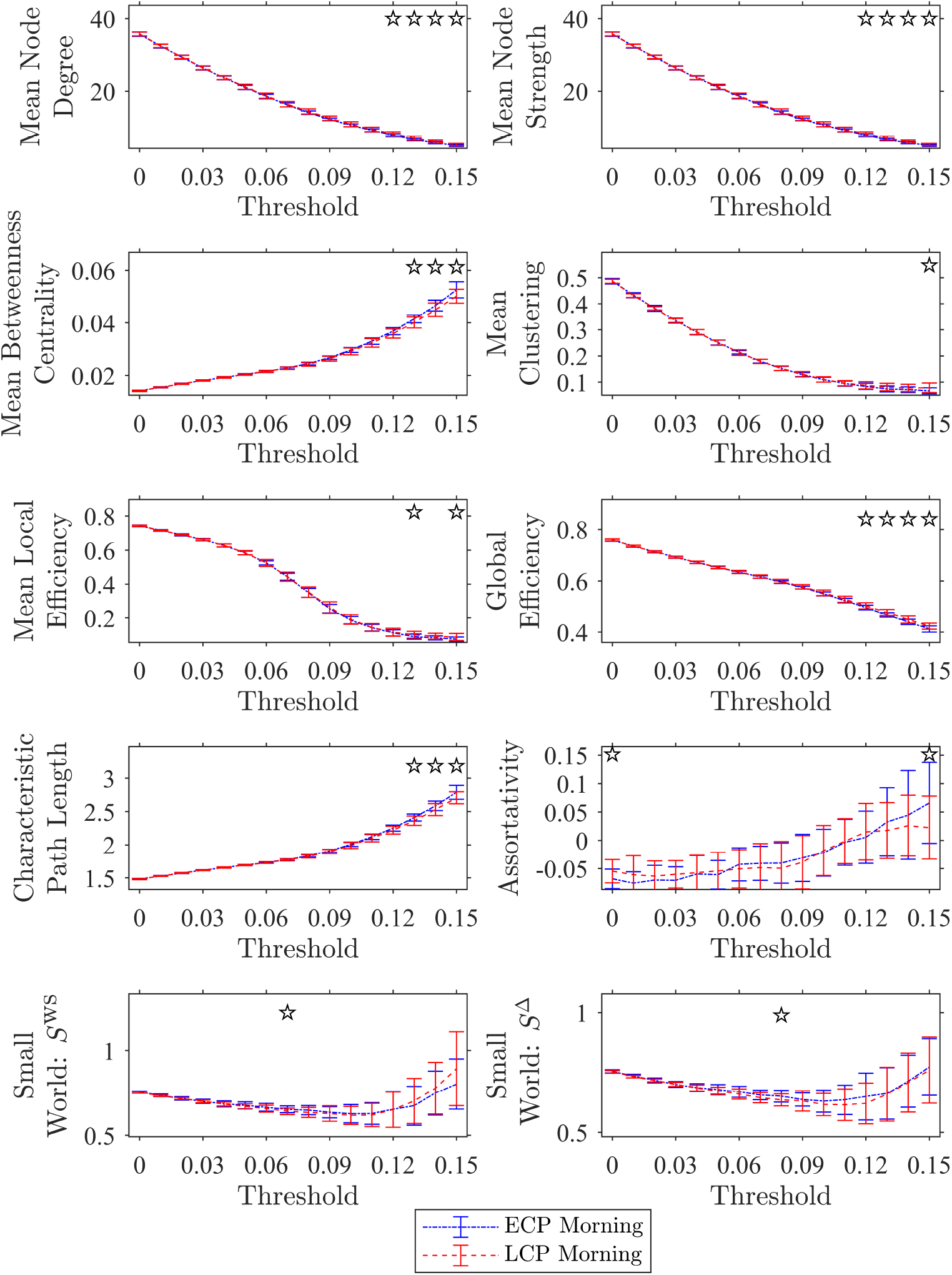
Distribution of Graph Metrics for the Morning Scan: Whole Network - Binarized. The values for various graph metrics for both ECPs and LCPs within the context of the whole network when binarized for non-negative thresholds increasing in step of 0.01 until the percolation threshold for the morning scan. The standard deviation between participants in that group is shown by the error bars. The ✩ indicates where the difference between ECPs and LCPs is significant *p* < 0.05 (uncorrected).

**Figure 11:**
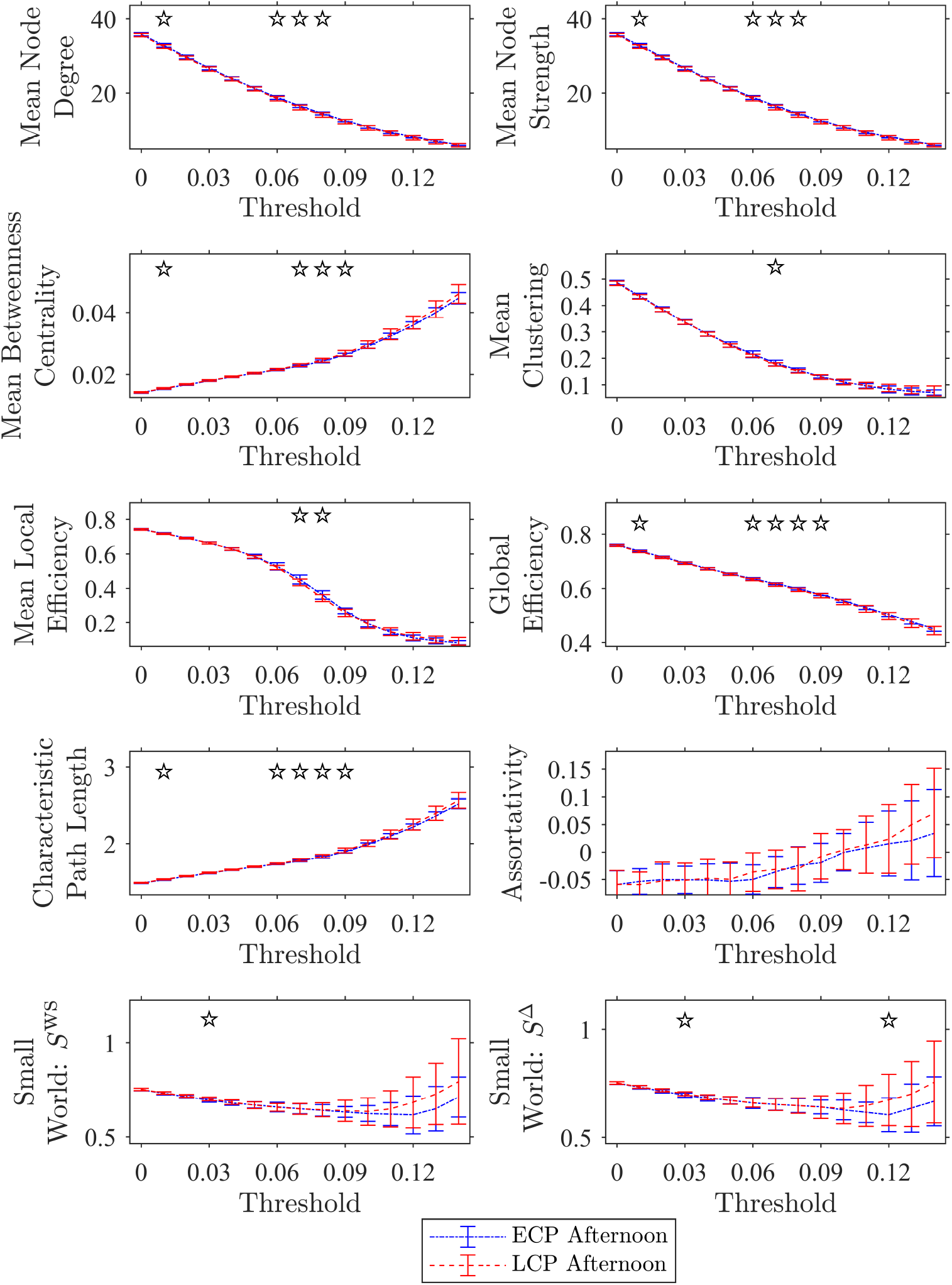
Distribution of Graph Metrics for the Afternoon Scan: Whole Network - Binarized. The values for various graph metrics for both ECPs and LCPs within the context of the whole network when binarized for non-negative thresholds increasing in step of 0.01 until the percolation threshold for the afternoon scan. The standard deviation between participants in that group is shown by the error bars. The ✩ indicates where the difference between ECPs and LCPs is significant *p* < 0.05 (uncorrected).

**Figure 12:**
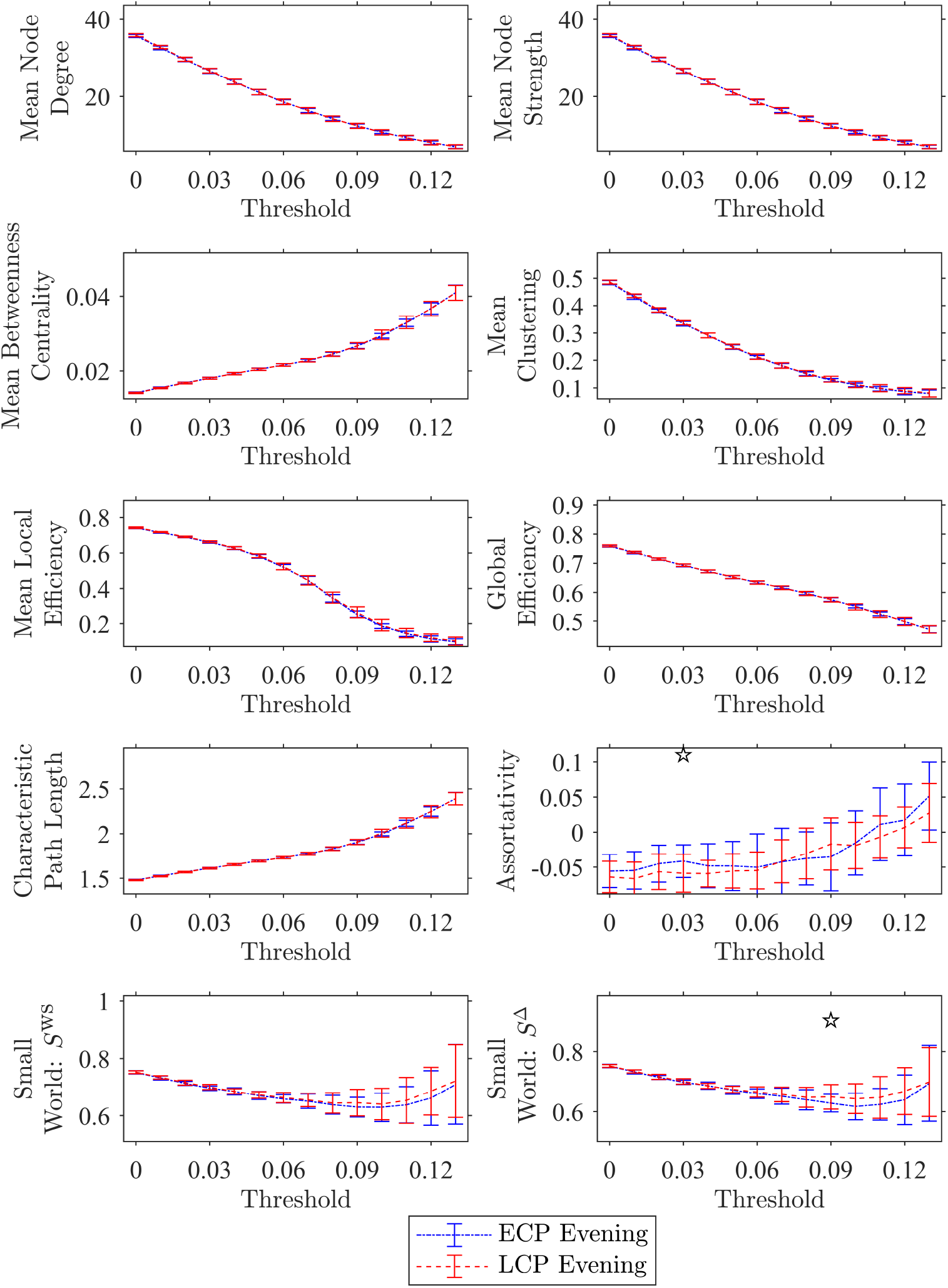
Distribution of Graph Metrics for the Evening Scan: Whole Network - Binarized. The values for various graph metrics for both ECPs and LCPs within the context of the whole network when binarized for non-negative thresholds increasing in step of 0.01 until the percolation threshold for the evening scan. The standard deviation between participants in that group is shown by the error bars. The ✩ indicates where the difference between ECPs and LCPs is significant *p* < 0.05 (uncorrected).

Figures 10 to 12 show that while there are some combinations of threshold, scanning session and graph metric that are significant with *p* < 0.05 this was only when uncorrected. Correcting for multiple comparisons occurring from testing at multiple thresholds as well as using different graph metrics using a Bonferroni or the less conservative Benjamini-Hochberg correction results in no significance. These results are consistent with those found independently in (Farahani, Fafrowicz, Karwowski, Bohaterewicz, et al. 2021).

## D Metadata

### D.1 Subjects Metadata

All the subjects were screened as ECPs or LCPs through a combination of actigraphy data, DLMO and CAR concentrations as well as their mid point of sleep on free days (MSF) when corrected for sleep debt (MSF_SC_). Therefore, subject metadata was considered to see if this offers an explanation for why the classifier is sensitive to certain subjects.

The metadata does seem to offer some explanation for why the classifier is so sensitive to the removal of certain subjects, especially in the case of ECPs as the MSF and DLMO appear to be lower for the subjects who were incorrectly labelled in the Afternoon and Evening scan. However, when corrected for multiple comparisons using Benjamini-Hochberg test no significance remains.

Therefore, across the three scanning sessions there no significant information to support that the metadata of subjects provides explains the sensitivity of the classifier to a subjects removal especially. This is further supported because, across the scanning sessions there is no consistency in the subjects whose removal has the biggest effect. Indeed, across all 3 scanning sessions the removal of 21 subjects leads to a high difference in the accuracy. Therefore, over half of the subjects cannot be considered outliers according to the metadata or the data would be fundamentally flawed.

